# Submersion and oxidative stress triggers pyrenoid formation, carbon-concentration-related protein remodeling and sub-plastidial rearrangements in hornworts

**DOI:** 10.1101/2025.05.05.652161

**Authors:** Svenja I. Nötzold, Susan Hawat, Thomas Stach, Stéphanie Ruaud, Péter Szövényi, Michael Hippler, Susann Wicke

## Abstract

Plastids are important for controlling acclimation of plants to environmental changes. In hornworts, chloroplasts may contain a RuBisCO-enriched protein matrix, a pyrenoid-like structure, which enables them to perform a biophysical carbon concentration mechanism (CCM) at the single-cell level– a unique feature among land plants. However, much remains unknown about the function, formation, and regulation of hornwort pyrenoids, especially as they are unaffected by changes in atmospheric CO_2_. Here, we tested whether submersion and hyperoxia induce pyrenoid formation and CCM. By subjecting *Anthoceros agrestis*, a pyrenoid-forming hornwort species, and *A. fusiformis*, which develops no pyrenoids, to a series of submersion experiments and analyzing their molecular, physiological, and cell-morphological response patterns using label-free proteomics and transmission electron microscopy, with additional *in silico* analysis, we identified a core set of CCM candidate genes. Under submersion, both species expressed CCM-associated protein homologs, whereas hyperoxia induced or diminished the expression of CCM-like homologs in a species-specific manner. We discovered that a carbonic anhydrase, a CAH3 homolog, as well as thylakoid bicarbonate transporter (LCI11-like) are upregulated under both conditions in *A. agrestis*, but not in *A. fusiformis*, suggesting that an algae-like mechanism of bicarbonate pumping into the thylakoid lumen and CO_2_ conversion exists in *A. agrestis*. Corroborating these molecular findings, an ultrastructural analysis of plastids revealed increases in pyrenoid-like structures and rearrangements during submersion in *A. agrestis*, whereas *A. fusiformis* accumulated lipid droplets between thylakoid stacks. Together, our data highlight hornworts’ distinct acclimation strategies to adverse environmental conditions, highlighting the relevance of their pyrenoids and CCM.

**Significance statement:** Carbon concentration via pyrenoids, densely packed matrices of a CO_2_-fixing enzyme, holds significant biotechnological promise for enhancing crop resilience. Studying mechanisms underlying pyrenoid formation in hornworts—the only land plants to naturally develop pyrenoids—is of exceptional interest due to their evolutionary closeness to crops. This study closes a knowledge gap by showing that hornworts use pyrenoids and a carbon-concentrating mechanism to adapt to submersion and hyperoxia, allowing them to thrive under regular transitions between atmospheric and submerged environments, resembling conditions during plants’ water-to-land transition. Species lacking pyrenoids accumulate lipid droplets instead, in addition to altering developmental and physiological pathways. Together, this work highlights hornworts’ remarkable versatility and untapped potential in advancing our understanding of plant adaptation to terrestrial life.

## Introduction

Photosynthetic organisms have independently evolved intricate mechanisms to enhance the efficiency of carbon fixation multiple times, primarily through Carbon Concentrating Mechanisms (CCMs). These adaptations address the limitations posed by ribulose-1,5-bisphosphate carboxylase/oxygenase (RuBisCO), the world’s most abundant enzyme responsible for carbon fixation, which has a relatively low affinity for CO_2_ and can also react with O_2_, leading to photorespiration and reduced photosynthetic efficiency. The evolution of CCMs represents a critical adaptive response to environmental pressures and photosynthesis’ innate biochemical limitations, allowing plants to thrive in varied and challenging habitats.

CCMs have evolved in various lineages, including cyanobacteria, algae, and vascular plants (1). In cyanobacteria and some algae, CCMs involve the active transport of inorganic carbon (CO_2_ and bicarbonate, HCO_3_^−^) into the cells, concentrating it around RuBisCO. Vascular plants, especially in arid and high-temperature environments, use C_4_ and CAM photosynthesis. C_4_ plants, like maize and sugarcane, spatially separate CO_2_ fixation and the Calvin cycle by means of a four-carbon compound and specialized leaf cells. This adaptation significantly reduces photorespiration and allows the plant to thrive under conditions of intense light and heat. CAM plants, such as cacti, temporally separate CO_2_ uptake and fixation, with nighttime CO_2_ storage and daytime usage, enabling efficient photosynthesis in extremely arid environments. In contrast, algae and a single land plant lineage—hornworts—use highly specialized structures known as pyrenoids, where RuBisCO is densely packed, to minimize its exposure to O_2_ and enhance CO_2_ fixation efficiency (2).

Pyrenoids enable a biophysical carbon-concentrating mechanism (CCM), best studied in the green alga *Chlamydomonas reinhardtii* (3). Such biophysical CCM relies on four crucial components for its operation ((2, 4–10), Table 1): Initially, inorganic carbon (HCO_3_^−^) is actively transported into the cytosol and plastids through HLA3 and LCIA transporters located in the plasma membrane and outer envelope of the plastid, respectively. Inside the plastid stroma, HCO_3_^−^ moves into the thylakoid channels (BST1-3, renamed LCI11A-C) and undergoes conversion into CO_2_ inside the thylakoid lumen by CAH3, powered by an acidic luminal pH that results from light-driven electron transfer. In the subsequent stage, the pyrenoid matrix plays a vital role, mainly consisting of RuBisCO associated with particular proteins, including the specific linker protein EPYC1 (2, 11). Ultimately, a CO_2_ leakage barrier forms due to starch organization guided by proteins like SAGA1 and SAGA2, and surrounding thylakoid membranes form thylakoid channels. CO_2_ that escapes the pyrenoid will be retransformed to HCO_3_^−^ via carbonic anhydrases (LCIB/LCIC), supported by an alkaline stromal pH generated by photosynthetic electron transfer.

**Table 1.**
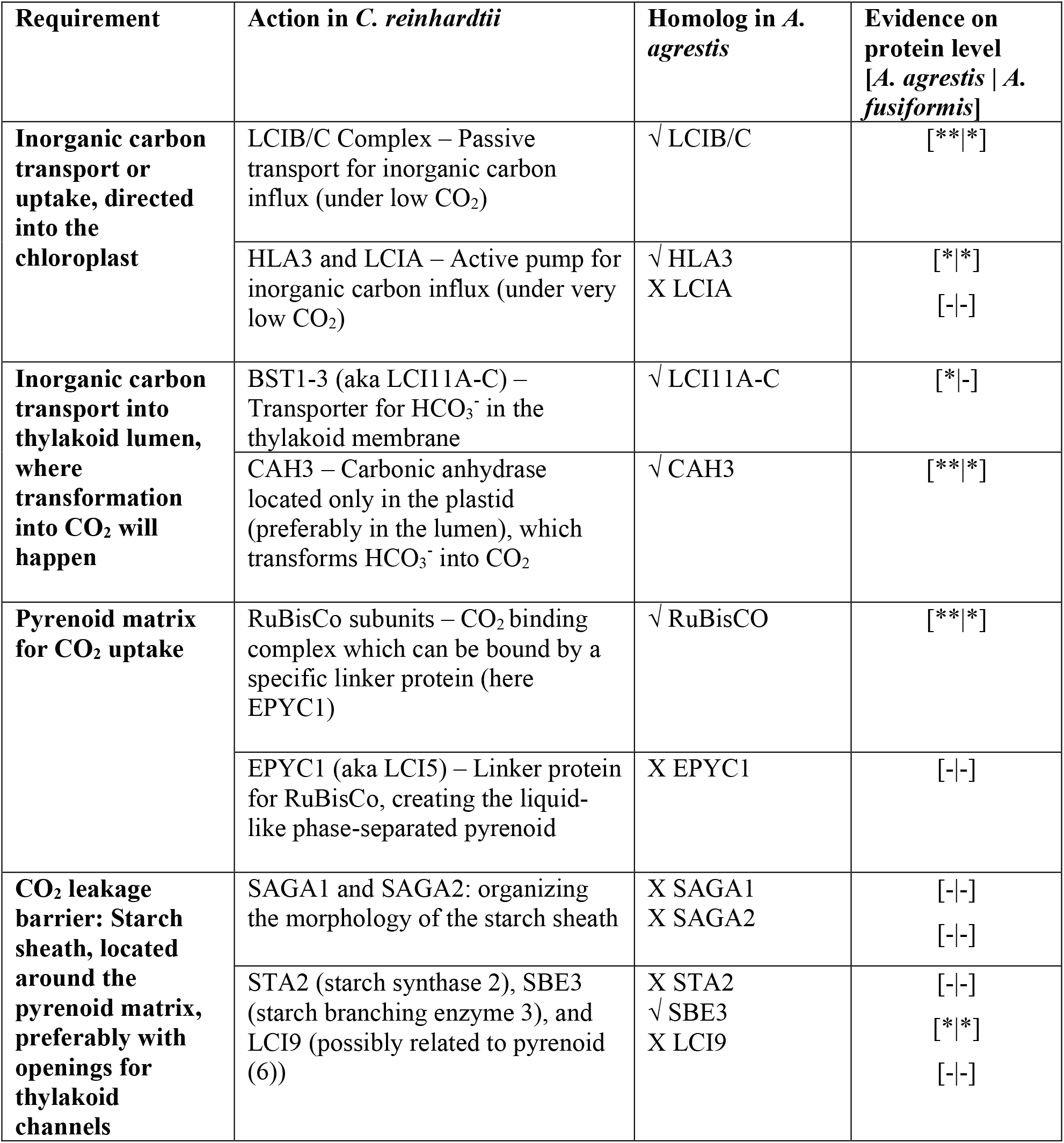
Minimal requirements for a pyrenoid-based biophysical CCM: Requirements and process description with involved proteins are based on *C. reinhardtii* studies and crop improvement by a pyrenoid-based CCM. Homologs of the mentioned important CCM-involved proteins in *A. agrestis* are indicated with presence by a check symbol while missing homologs are specified with a cross. Evidence on protein level, by identification through LC-MS/MS, is indicated with * (present) or – (missing), if presence in both species was confirmed, the number of * indicates the expression difference between both species, with ** representing higher expression than *.

Gene homology analysis suggests a certain degree of genetic conservation between the pyrenoid-based CCM in hornworts and that of algae (12). However, in hornworts, CCM associated with pyrenoid-like structures exhibits significant differences compared to that observed in algae, with multiple independent instances of pyrenoid gain and loss across various species (13). The presence or absence of pyrenoids does not seem to hinge on the type of plastid; rather, it appears to depend on other factors such as the position of the cell within the gametophyte or sporophyte (14). Pyrenoid-like structures in hornworts do not necessarily conform to a uniform morphology, often appearing as multiple pyrenoids instead of a single one (15). Although hornwort pyrenoids contain RuBisCO, the characteristic thylakoid channels involved in CO_2_ transformation and transportation into the pyrenoid matrix appear to be absent (16), whereby recent studies on hornwort CCM-related protein localization identified algae thylakoid channel typical proteins in the thylakoid stacks surrounding the pyrenoid area in hornworts (17, 18). Starch granules surround pyrenoid-like structures, although they may also occur near the chloroplast envelope. Interestingly, there is evidence of low C_4_-like organic matter carbon isotope discrimination in pyrenoid-containing hornworts, comparable to those seen in C_3_ plants (19). Nevertheless, much remains unclear about the specific functions and mechanisms underlying hornwort CCM.

Introducing a pyrenoid-based CCM into crop plants carries substantial biotechnological promise. In particular, gaining insights into pyrenoid formation and CCM regulatory mechanisms in hornworts may offer a tremendous advantage in engineering biophysical CCMs in crops. This potential advancement is due to hornworts’ closer evolutionary relationship with angiosperms compared to algae (20, 21), making them better models for cross-species genetic manipulation. Several hornwort species possess plastids with pyrenoid-like structures as observed in algae, while also exhibiting land plant features like a cuticle and stomata (22, 23). Pyrenoids are a unique feature among hornworts as land plants. In the *Dendroceros* genus, all species form pyrenoids, while genera such as *Leiosporoceros, Paraphymatoceros, Megaceros*, and *Phaeomegaceros* lack them entirely. Other genera, including *Anthoceros, Folioceros, Phaeoceros, Notothylas, Phymatoceros*, and *Nothoceros*, contain species with and without pyrenoids (14, 24, 25). The shape of pyrenoids in hornworts is highly diverse, but immunogold labeling has consistently shown that they accumulate RuBisCO in their matrix (15). Therefore, understanding hornwort biology and its specific adaptation to environmental challenges will offer valuable insights into plant adaptation to terrestrial life (26). On the other hand, harnessing this evolutionary link might enable us to effectively adapt and implement CCM strategies of hornworts to enhance crop plant resilience and productivity in harsh conditions or photorespiratory stress.

CO_2_ availability plays an important role in the CCM, often determining the path of the process. For example, gas diffusion in water is around ten thousand times slower than in air, while CO_2_ is not as abundant as other inorganic carbon sources (27). Therefore, concentrating carbon and turning it into CO_2_ to hinder the fixation of O_2_ is often an adaptation to an aquatic environment. However, in *C. reinhardtii*, pyrenoid-based CCM also depends on night and day rhythm (28) or hyperoxia (29). In contrast, the environmental parameters that induce CCM and pyrenoid formation in hornworts remain unclear, and the pattern of pyrenoid presence across species is puzzling. The relationship between atmospheric CO_2_ levels and the gain or loss of hornwort pyrenoids remains unresolved: although earlier work found no clear association (13), more recent phylogenetic studies suggest otherwise (30). Adding to the complexity, low CO_2_ conditions fail to induce a transcriptional response in hornwort CCM. (17) Above that, several aquatic hornwort species possess no pyrenoids (31), whereas species with pyrenoids show higher dissolved inorganic carbon pools compared to species without pyrenoids (32). Collectively, these findings not only challenge a simple CO_2_-driven regulation model but also highlight unresolved questions about the evolutionary pressures and environmental factors shaping the presence and function of CCM in hornworts.

Hornworts thrive in both aquatic and terrestrial habitats, where they commonly experience regular transitions from atmospheric to submersed conditions with periods of light fluctuation, temperature variation, dryness, and, therefore, changes in CO_2_ availability, potentially resembling conditions during plants’ water-to-land transition a half billion years ago. Therefore, the hornwort CCM may be a unique adaptation to periodic flooding—a hypothesis we challenge in this study. To this end, we utilize a comparative label-free proteomics approach to investigate the effects of two-day submersion and hyperoxic conditions through H_2_O_2_ treatment on the formation of pyrenoid-like structures and induction of the CCM in two hornwort species: *Anthoceros agrestis*, which forms pyrenoids, and *A. fusiformis*, which does not. Ultrastructural analysis using transmission electron microscopy of both species’ plastids under submersion and hyperoxia reveals species-specific cell-morphological changes, supporting the primary molecular findings from our tandem mass spectrometric analyses. Additionally, mining our untargeted protein data alongside protein homology analysis of known algal CCM proteins enabled us to identify a core set of functional protein players in hornwort CCM and hornworts’ response strategies to adverse environmental conditions.

## Results

### Global protein identification in hornworts

Label-free proteomic analysis of samples from the two hornwort species *A. agrestis* and *A. fusiformis* collectively identified 2,592 proteins with high confidence (p<0.01). Among these, 2,214 proteins (85%) were detected in *A. agrestis* samples, while 1,977 proteins (76%) were identified in *A. fusiformis* samples. A total of 1,599 proteins (61%) were expressed in both hornwort species, whereas 615 proteins (23%) were exclusive to *A. agrestis* and 379 proteins (14%) to *A. fusiformis* (Supplemental Figure S1). An overrepresentation analysis of Gene Ontology terms (GO-ORA) for all identified proteins in *A. agrestis* and *A. fusiformis* separately revealed that both hornwort species expressed proteins involved in photosynthesis and the generation of precursor metabolites and energy. These processes included energy derivation through the oxidation of organic compounds, the glucose catabolic process, and the electron transport chain, with most of these proteins located in the thylakoid lumen (Supplemental Figure S2A). In addition to these proteins, proteins related to responses to cold and bacterial challenges were highly prevalent, indicating that the submersion treatment regulates similar abiotic and biotic response pathways. At the molecular level, both species exhibited significant involvement in replication and transcription, particularly through purine nucleotide and ribonucleotide binding. Most of these proteins were located in the cytosol or associated with plastoglobules.

### Proteins with species-exclusive occurrence

Proteins exclusively expressed in *A. agrestis* were involved in photosynthesis but predominantly in translational processes associated with ribosomes and present in the cytosol (Supplemental Figure S2B). The molecular functions of these *A. agrestis*-specific proteins included tetrapyrrole binding, which is crucial for the formation of chlorophyll. Additionally, ligase activity and macrolide binding were also noted. Macrolides, known to block protein synthesis by interacting with the large ribosomal subunit, appear to bind in *A. agrestis*, potentially facilitating higher rates of protein translation. In contrast, proteins exclusive to *A. fusiformis* were implicated in cellular responses to heat, small molecule catabolic processes, and, intriguingly, spore germination and regulation (Supplemental Figure S2C). Furthermore, phytohormone-related ABA degradation, isomerase activity, and hydrolase activity for C-N bonds (excluding peptide bonds) and bonds between sugars were present in *A. fusiformis*. These proteins were located in the mitochondrial membranes, peroxisomes, early endosome membranes, and P-bodies, indicating processes related to photorespiration and reduced translation through mRNA degradation.

### Changes in protein expression during submersion

Principal component analysis (PCA) of 48 hornwort samples subjected to submersion in either H_2_O or H_2_O_2_ revealed clustering by species and treatment (Supplemental Figure S3). *A. agrestis* samples submerged in H_2_O for 24 hours (AH24) were closer to the control than those submerged for 48 hours (AH48), showing higher variance. A total of 1,384 proteins (62% of identified proteins) were significantly differentially expressed between treatments in *A. agrestis*, while 1,309 proteins (66%) were identified in *A. fusiformis* (p<0.05), including proteins only present in one treatment. When considering only proteins present across all treatments, 432 proteins were significantly expressed in *A. agrestis* and 436 in *A. fusiformis*. Of these, 126 proteins were common to both species, with the rest almost equally split between the species (Supplemental Figure S4). Significant changes in these common proteins were observed in those involved in disaccharide processes, chloroplast targeting, and nucleotide binding, suggesting transcriptional upregulation of plastid-targeted proteins (Supplemental Figure S5A). Proteins associated with cold response also reappeared, indicating a shared response between species. *A. agrestis* showed highly expressed photosynthesis-related proteins between treatments, primarily in the cytosol and ribosomes, indicating roles in protein translation and chloroplast transport (Supplemental Figure S5B). *A. fusiformis* displayed numerous proteins linked to cold response, carbohydrate catabolic processes, and photosynthesis, although fewer than *A. agrestis* (Supplemental Figure S5C). These proteins were located in the thylakoid lumen, stromule, and associated with plastoglobule activity. In *A. agrestis*, the Top 20 significantly expressed proteins were mainly involved in DNA replication, RNA ligase activity, mRNA binding, and tryptophan biosynthesis, suggesting roles in transcription and translation (Supplemental Figure S6A). In contrast, *A. fusiformis* proteins were involved in nuclear processes, reproductive development, and the auxin pathway, including sugar transport and terpene synthesis, possibly related to steroid or oil production (Supplemental Figure S6B). Together, these results show that both species exhibit significant protein expression changes in response to submersion, though with distinct focuses: *A. agrestis* modulates photosynthesis, and *A. fusiformis* activates developmental processes and auxin pathways.

### Time- and treatment-specific protein clusters

We performed weighted correlation network analysis (WGCNA) separately for both hornwort species, identifying 13 modules in *A. agrestis* and 16 in *A. fusiformis* (Figure 1). Among the three unique responses of *A. agrestis* was an upregulation of proteins associated with reproductive development during submersion in water and downregulation in hyperoxic conditions (tan moduleA, H_2_O ↑, H_2_O_2_ ↓), a downregulation of mitotic and plastid processes in water after 24 hours and consistently under hyperoxia (yellow moduleA, H_2_O 24h & H_2_O_2_ ↓), as well as a downregulation of proteins related to sugar response in water for the first 24 hours, and even more so after 48 hours and under hyperoxia (black moduleA; H_2_O 24h ↓, H_2_O 48h & H_2_O_2_ 48h ↓↓). In contrast,

**Figure 1.**
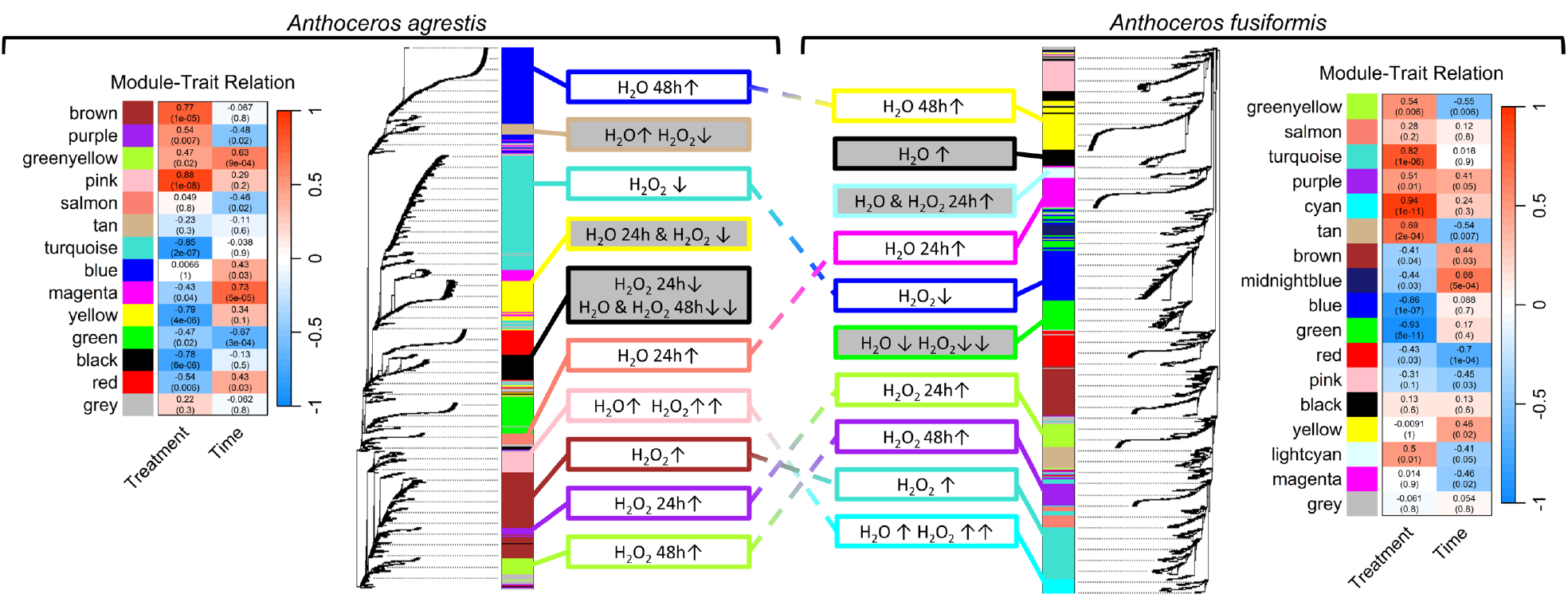
Weighted correlated network analysis (WGCNA) of identified proteins from LC-MS/MS: Separately calculated protein expression networks for both species. Grey-filled expression boxes indicate unique modules for the corresponding species, while colored lines connect two modules with the same expression profile between species. *A. agrestis* was identified with 13 modules of which 10 were submersion-relevant. *A. fusiformis* was identified with 16 modules and 10 submersion-relevant modules. The expression boxes describe the down- or up-regulation of proteins within the module and their behavior among the submersion treatments. An up arrow indicates upregulation, while a down arrow indicates downregulation, with two arrows indicating stronger expression than just one. Next to each dendrogram is the module-trait relationship shown, given the correlation and p-value (in parenthesis) for the corresponding module and time or treatment as tested traits.

*A. fusiformis* responded with photosynthesis downregulation in both conditions (green moduleF, H_2_O ↓, H_2_O_2_ ↓↓), an upregulation of peroxisome activity under water and during the first 24 hours of hyperoxia (lightcyan moduleF, H_2_O & H_2_O_2_ 24h ↑), and an upregulation of cold response proteins during water submersion (black moduleF, H_2_O ↑). A module-trait relationship analysis indicated that *A. agrestis* predominantly activates proteins involved in photosynthesis and light response under both submersion and hyperoxic conditions (pink moduleA, H_2_O ↑, H_2_O_2_ ↑↑; p << 0.001, corr = 0.88), whereas *A. fusiformis*’ responses specifically with the expression of proteins located in mitochondria as well as those associated with nutrient-reservoir activity and photorespiration (cyan moduleF, H_2_O ↑, H_2_O_2_ ↑↑; p = 1e-11, corr = 0.94). Interestingly, despite similar expression profiles, the pink moduleA of *A. agrestis* and the cyan moduleF of *A. fusiformis* displayed differences in the nature of their functional proteins. *A. agrestis* upregulated CO_2_ fixation and photosynthesis under submersion, with an even greater increase during hyperoxia. In contrast, *A. fusiformis* shifted towards mitochondrial photorespiration under the same conditions. While both species shared expression profiles, the predicted protein functions varied based on GO-term overrepresentation. Notably, proteins homologous to known algal CCM proteins were not confined to specific modules but were distributed across different expression profiles (Figure 4, Supplemental Table S1). In addition to co-expression network analysis, we found exclusive proteins induced solely during submersion treatments (Figure 2, n = 304). The highest number of these treatment-exclusive proteins was observed in *A. agrestis* after 48 hours of submersion (n = 79), followed by *A. fusiformis* after 24 hours underwater (n = 27). Specifically, the 24-hour water submersion in *A. agrestis* coincided with pyrenoid induction, revealing 10 proteins involved in immune response and amino acid activation, but none were CCM-related homologs. Candidate proteins involved in pyrenoid formation may either be exclusively present at the induction time point (first 24 hours of submersion) or already present in control treatments at lower expression levels compared to the 24-hour submersion treatment. Together, these findings underscore the species-specific responses and core CCM-like protein regulation in hornworts under various submersion treatments. The pyrenoid-forming *A. agrestis* primarily enhances photosynthetic processes and CO_2_ fixation, while the pyrenoid-lacking *A. fusiformis* shifts towards developmental and photorespiration pathways. This reflects distinct adaptive strategies to submersion stress in each species.

**Figure 2.**
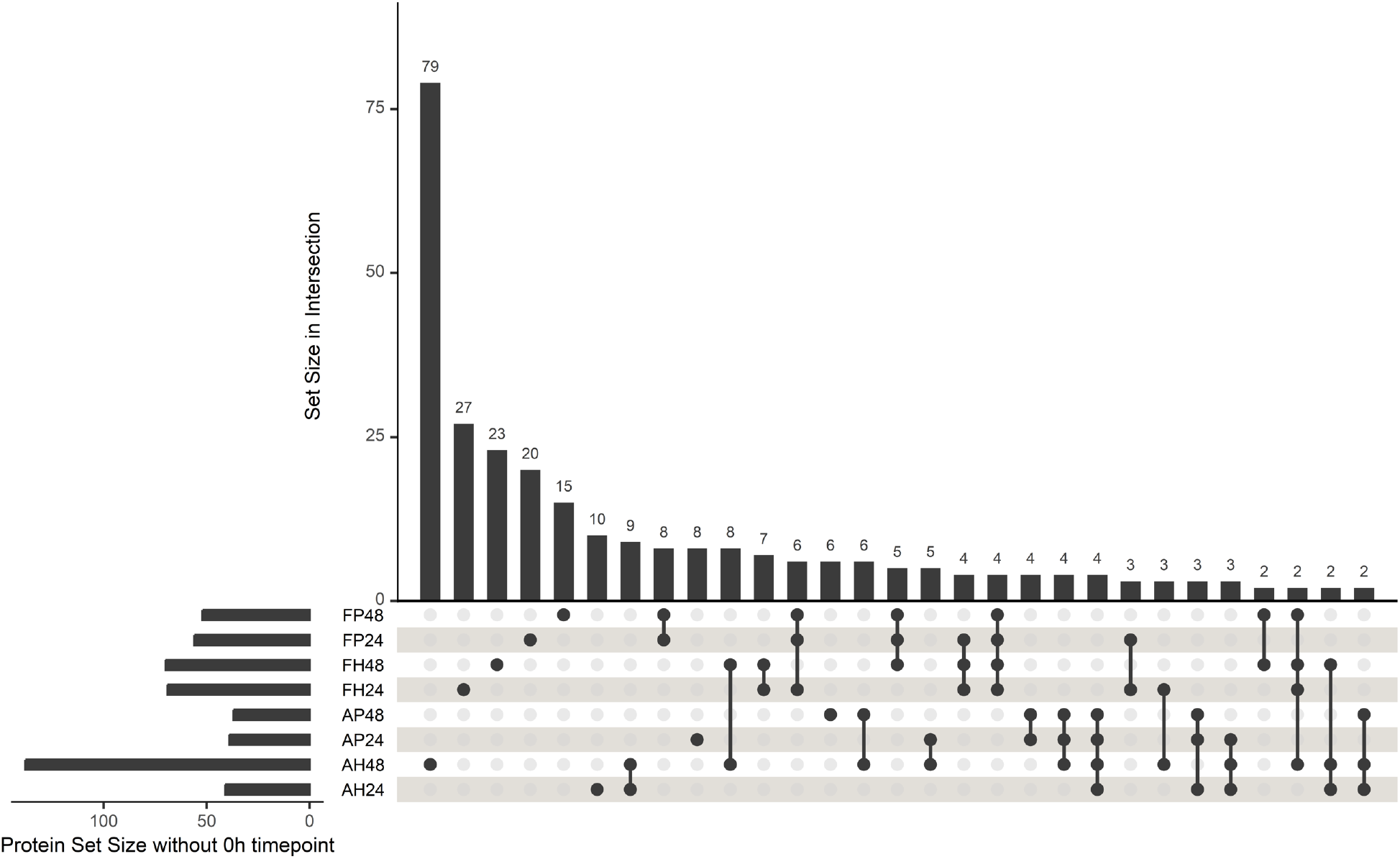
UpSet plot for proteins only present under submersion: Groups are named by treatment and the total protein set size is given. Protein set sizes in intersections are in descending order with the total size given by the number above each bar. Intersections are given by filled circles and intersections of multiple groups are indicated by connection trough lines. Only proteins were considered, that were present in any submersion treatment, but not in any Control group, a total of 304 proteins were distributed among all intersections. Not all intersections are shown, intersections with 0 or 1 member are not displayed.

### Inner-plastidial rearrangements during submersion over time

Analysis by transmission electron microscopy (TEM) revealed a significant increase in the number of pyrenoid-like structures in *A. agrestis* after the first 24 hours of water submersion, while *A. fusiformis* showed no such structures under any condition (Figure 3). In *A. agrestis*, pyrenoid-like structures in normal conditions are primarily located in the middle of the plastid, with starch fields spreading across the plastid but not necessarily adjacent to the pyrenoid, unlike in algae. Lipid droplets (LDs) are dispersed within the plastid. In contrast, *A. fusiformis* plastids feature numerous LDs grouped in the stroma between thick thylakoid stacks. After 24 hours of submersion, *A. agrestis* shows a significant increase in pyrenoid-like structures (p < 0.001), which cluster centrally in the plastids. In *A. fusiformis*, LDs enlarge and vary in size, forming large groups or multiple smaller clusters, with plastids becoming more rounded and swollen. After 48 hours of submersion, *A. agrestis* displays swollen LD groups forming long strands, while the number of pyrenoid-like structures decreases. *A. fusiformis* exhibits increased open stroma spaces between thylakoids, partially filled with LDs. Under hyperoxia for 24 hours, *A. agrestis* shows a decrease in pyrenoids with an increase in starch field size, whereas *A. fusiformis* responds similarly to its submersion response, but also forms extensive thylakoid strings with sparse LD groups. After 48 hours of hyperoxia, *A. agrestis* occasionally produces pyrenoids, and *A. fusiformis* shows two plastid morphologies: enlarged plastids with large starch fields and small LD groups, or small plastids with few starch fields, large variable-sized LD groups, and disorganized thylakoids. Together, these results show that submersion significantly induces pyrenoid formation and cell-morphological changes in *A. agrestis* within the first 24 hours, while hyperoxia has minimal effects on pyrenoid formation, and both submersion and hyperoxia affect LD organization within 48 hours in this species. In contrast, *A. fusiformis* responds to submersion and hyperoxia primarily through LD formation and relocation, with hyperoxia causing plastids to exhibit two distinct phenotypes, potentially indicating functional plastid differentiation during adverse conditions.

**Figure 3.**
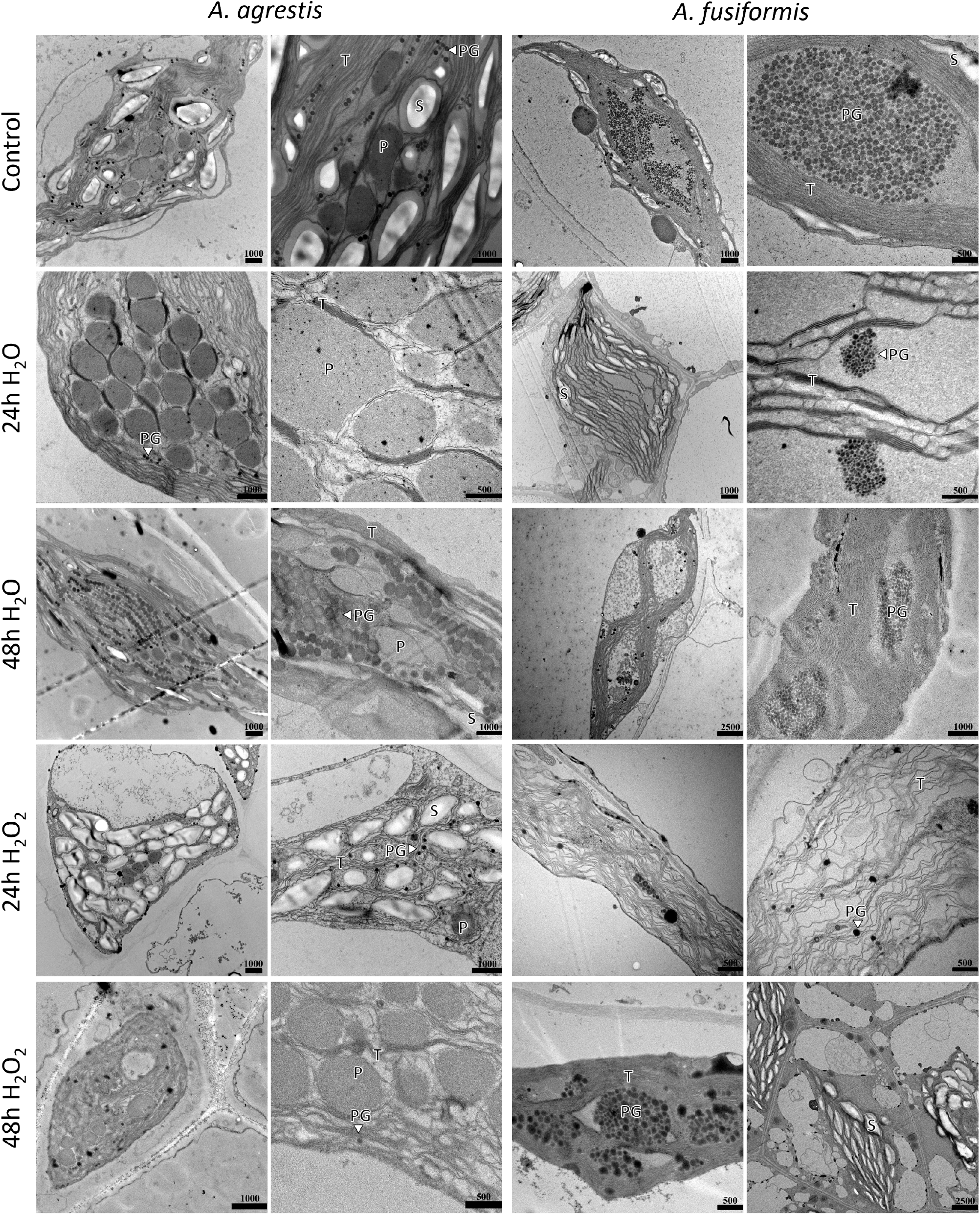
Transmission Electron Microscopy (TEM) of hornworts during submersion: Both hornworts *A. agrestis* (with pyrenoid) and *A. fusiformis* (without pyrenoid) were sampled for control without submersion and for 24h and 48h under submersion with water (24 H_2_O and 48 H_2_O) and water with additional hydrogen peroxide (24 H_2_O_2_ and 48 H_2_O_2_). Abbreviations: Pyrenoid (P), Thylakoid membranes (T), Starch fields (S), Lipid droplets/plastoglobules (PG). Scale bars, in nanometers (nm), are indicated on each picture. The full picture archive, used for the measurements and follow-up statistics, can be found in the supplement.

**Figure 4.**
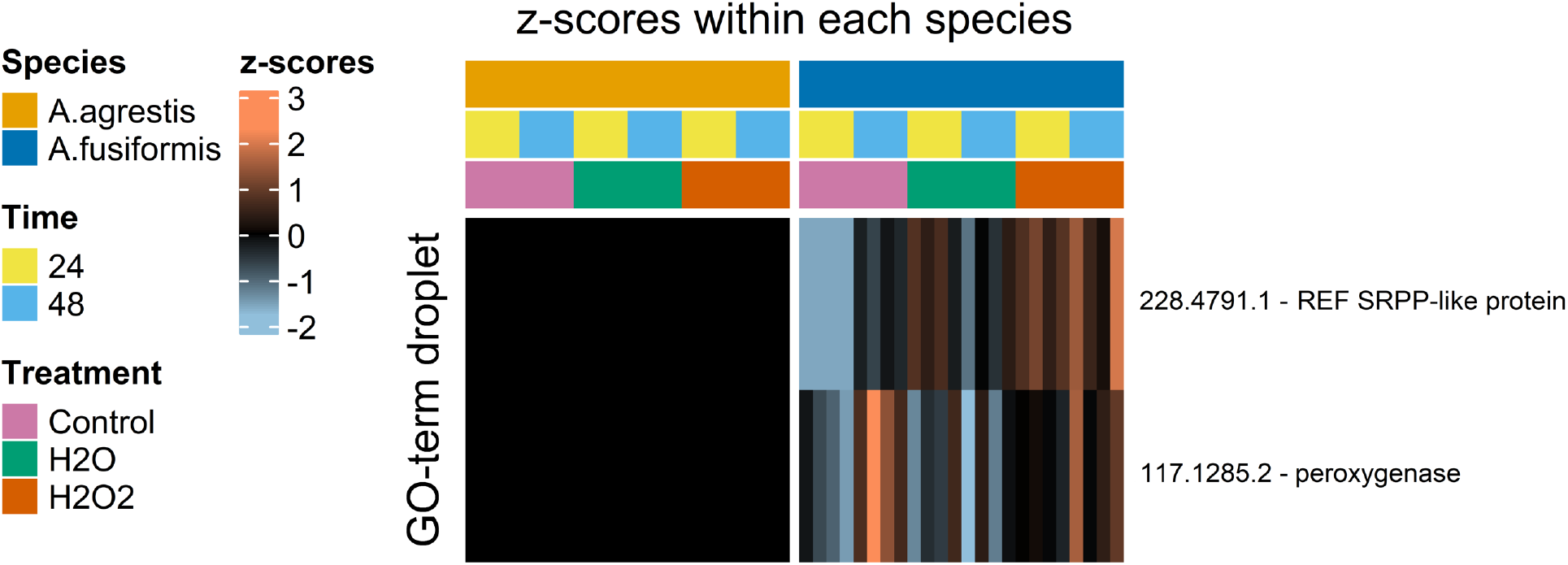
Heatmaps of “lipid droplet”-associated proteins in *A. agrestis* and *A. fusiformis*. Intensities are shown in z-scores, calculated between all groups of each species (left), and shown in log_2_ across both species (right). The annotation above the heatmap indicates the time point, treatment, and species of each group. *A. agrestis* gene IDs are displayed with the corresponding annotation; see methods for more information on the annotation. The two shown proteins were not detected in *A. agrestis* samples, only in *A. fusiformis*.

### Compositional changes in pyrenoids during submersion

Pyrenoid structures occurred under all treatments in *A. agrestis*. Analysis of 169 pyrenoids across all treatments revealed significant differences in the relative distribution of low- and high-density areas in transmission electron microscopy (TEM) images (Kruskal-Wallis test, p < 0.001). Under normal conditions, pyrenoids exhibited a predominance of low-density regions, covering approximately 60% of the pyrenoid area, whereas high-density regions accounted for 40% (Supplemental Table S10). However, exposure to oxidative stress for 48 hours significantly altered this distribution, with low-density regions decreasing to 55% and high-density regions increasing to 45%. Further pairwise comparisons confirmed that these shifts were statistically significant in *A. agrestis*. Notably, significant differences were observed between the 24-hour and 48-hour H_2_O_2_ treatment (Dunn’s test, p_adj._ < 0.001), as well as the 24-hour water submersion and 48-hour hyperoxia submersion (Dunn’s test, p_adj_ < 0.001), and control vs. 48-hour H_2_O_2_ submersion (Dunn’s test, p_adj_ < 0.001). These findings suggest that prolonged oxidative stress induces a measurable reduction in low-density regions and a corresponding increase in high-density areas, potentially reflecting protein condensation or phase transition changes within the pyrenoid. This structural reorganization may play a role in the regulation of CO_2_-concentrating mechanisms under stress conditions.

### Induction of subcellular structures during submersion

To assess CCM-related plastid rearrangements, we measured sub-cellular structures in *A. agrestis* plastids during pyrenoid formation (Supplemental Figure S7). Stacks of thylakoid membranes were commonly found between multiple pyrenoid-like structures, with significant differences observed only between control and submersion treatments. Thylakoid thickness was highest in the control (p = 0.01). The extension of the thylakoid lumen within the stacks around the pyrenoids decreased during submersion and hyperoxia (p < 0.001). Though less abundant in *A. agrestis* than in *A. fusiformis*, the mean diameter of lipid droplets (LDs) in *A. agrestis* decreased in the first 24 hours of water submersion, then increased after 48 hours compared to controls (p << 0.001). There was no significant difference over time in LD diameter under hyperoxia, but compared to control samples, significant differences were noted (24h H_2_O_2_ to Control, p <0.001; 48h H_2_O_2_ to Control, p = 0.005). The highest number of pyrenoid-like structures occurred after 24 hours of submersion and 48 hours of hyperoxia (p = 0.001), with the mean area of these structures being largest under control conditions (p < 0.001) and consistent across submersion treatments. The sum of pyrenoid-like structures per plastid (total area size) was highest after 24 hours of submersion, significantly higher than under hyperoxia (p = 0.03). These results show that *A. agrestis* has a higher number of pyrenoids per plastid after 24h submersion, without a corresponding increase in individual pyrenoid size. As a result, the total pyrenoid area—and thus the pyrenoid surface—per plastid is highest in *A*.*agrestis* while submersed due to the increased pyrenoid number. Interestingly, the smallest and largest pyrenoids were found under submersion (p = 0.024), with a significant difference between control and submersion at 24 hours (p = 0.019). These findings confirm that submersion triggers pyrenoid-like structure induction within the first 24 hours, accompanied by dynamic rearrangement of sub-organellar structures in the plastids of *A. agrestis*.

### Physiological performance during submersion and oxidative stress

We investigated the photosynthetic efficiency of the gametophytes of *A. agrestis* and *A. fusiformis* using Pulse-Amplitude Modulation (PAM) measurement, focusing on photosystem II (PSII) operating efficiency (Y(II)) and non-photochemical quenching (NPQ) under submersion (Supplemental Figure S8, Supplemental Table S7). In *A. agrestis*, Y(II) remained around 0.5 underwater over a 72-hour period, indicating stability in photosynthetic efficiency under normal submersion conditions. Similarly, *A. fusiformis* exhibited a Y(II) around 0.6 throughout the 72 hours, demonstrating that more than half of the absorbed quanta were converted into chemically fixed energy in both hornwort species, irrespective of submersion or recovery periods. Conversely, the hyperoxia negatively impacted photosynthetic efficiency compared to submersion in water. In *A. fusiformis*, Y(II) decreased to 0.4 after the first 24 hours of oxidative conditions in H_2_O_2_ but increased to 0.5 after an additional 24 hours. Upon removal of H_2_O_2_, Y(II) recovered to 0.6, suggesting a short-term adjustment of the photosynthetic apparatus under hyperoxic conditions. In contrast, *A. agrestis* experienced a significant drop in Y(II) to 0.2 after 24 hours (three-way ANOVA with Tukey’s HSD, p-value < 0.005; Supplemental Table S7) and 48 hours of hyperoxia (three-way ANOVA with Tukey’s HSD, p-value < 0.05; Supplemental Table S7). However, Y(II) increased to nearly 0.4 following the removal of H_2_O_2_ submersion, without significant difference from the time point before oxidative conditions. NPQ, representing the dissipation of excess light energy as heat, showed different trends under various conditions. Under standard conditions (0 hours), *A. agrestis* had a slightly lower NPQ (0.6) compared to *A. fusiformis* (0.7), whereby the difference in NPQ values increased under submersion. During submersion in water, NPQ values in *A. fusiformis* increased, while *A. agrestis* showed a reduction to 0.2, which later increased to 0.3 after 24 hours of recovery, indicating a decrease in quenching activity in line with a decrease in Y(II) in *A. agrestis* (n.s.). Under hyperoxia treatment, *A. agrestis* reduced NPQ to 0.4 (three-way ANOVA with Tukey’s HSD, p-value < 0.04; Supplemental Table S7), which increased to 0.6 during recovery. Similarly, *A. fusiformis* exhibited a drop in NPQ from 0.8 to 0.3 during H_2_O_2_ treatment (three-way ANOVA with Tukey’s HSD, p-value < 0.02; Supplemental Table S7), with recovery to 0.6 after 48 hours and further to 0.7 after submersion recovery. H_2_O_2_ treatment significantly decreased PSII efficiency in *A. agrestis*, also in the long run, whereas *A. fusiformis* demonstrated partial recovery after 24 hours without submersion. In contrast, submersion in water did not affect PSII efficiency, as Y(II) remained stable for both species. *A. agrestis* showed low NPQ, particularly under water submersion, while *A. fusiformis* maintained high NPQ over 72 hours. Although hyperoxia reduced NPQ in both species, recovery post-submersion nearly restored initial conditions.

### *A. fusiformis*-specific proteins for lipid droplet formation

Instead of the typical pyrenoid-like structures seen in *A. agrestis, A. fusiformis* accumulates LDs in analogous locations (Figure 3). A search for proteins with the GO-term “lipid droplet” identified two proteins expressed uniquely in *A. fusiformis* and undetected in *A. agrestis* (Figure 4). These proteins are present in most treatments and are particularly abundant during hyperoxia. Homology analysis with *C. reinhardtii* and *Arabidopsis thaliana* identified one of these proteins, *A. agrestis* protein 228.4791.1, as a caleosin. The caleosin family is typically associated with oil bodies or lipid droplets and plays a critical role in their structural integrity and function. Caleosins are also known to participate in the formation and maintenance of lipid droplets in the leaves of higher plants during periods of stress. For the gene product of *A. agrestis* protein 117.1285.2, no homolog was found in *C. reinhardtii*, but comparisons with the *A. thaliana* proteome suggest a similarity to a group of rubber elongation factor proteins, and the unstudied homolog might be newly implicated in a physiological response associated with submersion and hyperoxia. Together, these findings suggest that the LD response in *A. fusiformis* results from the specific activation of lipid biosynthesis and storage pathways.

### Differential regulation of CCM proteins

We predicted 768 CCM-like homologs in *A. agrestis*, of which 347 proteins were detected by LC-MS/MS. Of these, 269 showed significant expression differences between treatments in both species (p<0.05), constituting three main clusters per species (Supplemental Figure S9). In *A. fusiformis*, one cluster included proteins upregulated only during submersion, while the other two clusters showed downregulation during hyperoxia. The identified core subset of CCM-involved proteins in *A. fusiformis* was distributed across these three clusters, with HLA3 and CAH3 homologs downregulated under hyperoxia and LCIB/C and RBCS homologs upregulated during submersion. In the pyrenoid-present species *A. agrestis*, submersion predominantly upregulated the majority of significant CCM-like proteins, and some CCM-like proteins were also upregulated during hyperoxic conditions. Additionally, *A. agrestis* downregulated some CCM-like proteins in both treatments. The core CCM subset in *A. agrestis* was present in only two of the three clusters, with HLA3 homologs downregulated during submersion and LCI11 and RBCS homologs upregulated only under hyperoxia. Despite the absence of pyrenoids in *A. fusiformis*, it expressed CCM-like proteins, but with expression patterns different from those of *A. agrestis* (Supplemental Figure S9). These findings suggest that a form of CCM may operate in *A. fusiformis* despite the absence of pyrenoid-like structures characteristic of *A. agrestis*. Notably, *A. agrestis* did not exhibit consistent upregulation of CCM-related proteins across both submersion treatments, in contrast to the pyrenoid-lacking *A. fusiformis*. Differential expression analysis of CCM-like proteins revealed that submersion, CO_2_ limitation, and hyperoxia each affect *A. agrestis* in distinct ways, with minimal overlap in protein expression between treatments. In *A. fusiformis*, however, submersion in water resulted in expression patterns largely similar to the control, while hyperoxia induced a distinct expression profile.

### Expression of required core pyrenoid proteins

Considering the minimal requirements of a pyrenoid-based CCM, we defined a core set of CCM-like proteins (Supplemental Table S1) and analyzed their expression profiles in our hornwort species. A comparison of log_2_-transformed intensity values between *A. agrestis* and *A. fusiformis* revealed that most of these core CCM-like proteins are expressed at higher levels in *A. agrestis*. Notably, four of the core proteins are absent in *A. fusiformis*, and two are absent in *A. agrestis*. However, proteins such as LCIB/C and HLA3 exhibit similar expression rates in both species (Figure 5). Specific homologs of HLA3 and CAH3, as well as LCI11 and RBCS1, are induced under submersion and hyperoxia in *A. agrestis* but not in *A. fusiformis*. Additionally, a CAH3 homolog with a predicted plastid transit peptide (cTP) is present only during submersion in both species. Despite the presence of the core CCM-like protein subset in both species, their expression levels vary significantly under different treatments. In *A. fusiformis*, most core CCM-like proteins are not induced by hyperoxia and are often downregulated, with the exception of two HLA3 homologs that are upregulated. The cTP-containing CAH3 homolog is induced exclusively under water submersion, while another HLA3 homolog is induced by any treatment. Notably, key CCM-like proteins, such as LCI11 and multiple RBCS homologs, are absent in all *A. fusiformis* samples. In contrast, *A. agrestis* shows an induction of core CCM-like proteins under hyperoxia, similar to the induction of CCM in *C. reinhardtii* in response to reactive oxygen species (33). Homologs to LCIB/C, CAH3, and LCI11 are more highly expressed under hyperoxia compared to control and water submersion conditions. Except for the RuBisCO accumulation factor (RCA), the remaining core CCM-like proteins in *A. agrestis* are not downregulated under hyperoxia. Additionally, *A. agrestis* has core CCM-like proteins that appear unaffected by submersion, such as two RBCS homologs and an HLA3 homolog, a pattern not observed in *A. fusiformis*. The presence of pyrenoid-like structures in *A. agrestis* may influence the expression of specific core CCM-like proteins. For example, the presence of LCI11 in *A. agrestis* may be associated with its pyrenoid, while the absence of a pyrenoid in *A. fusiformis* might be linked to the non-expression of LCI11. In sum, these findings highlight the species-specific responses and the regulation of core CCM-like proteins under various environmental conditions, reflecting distinct adaptive strategies in *A. agrestis* and *A. fusiformis*, and emphasizing the potential influence of pyrenoid-like structures on protein expression.

**Figure 5.**
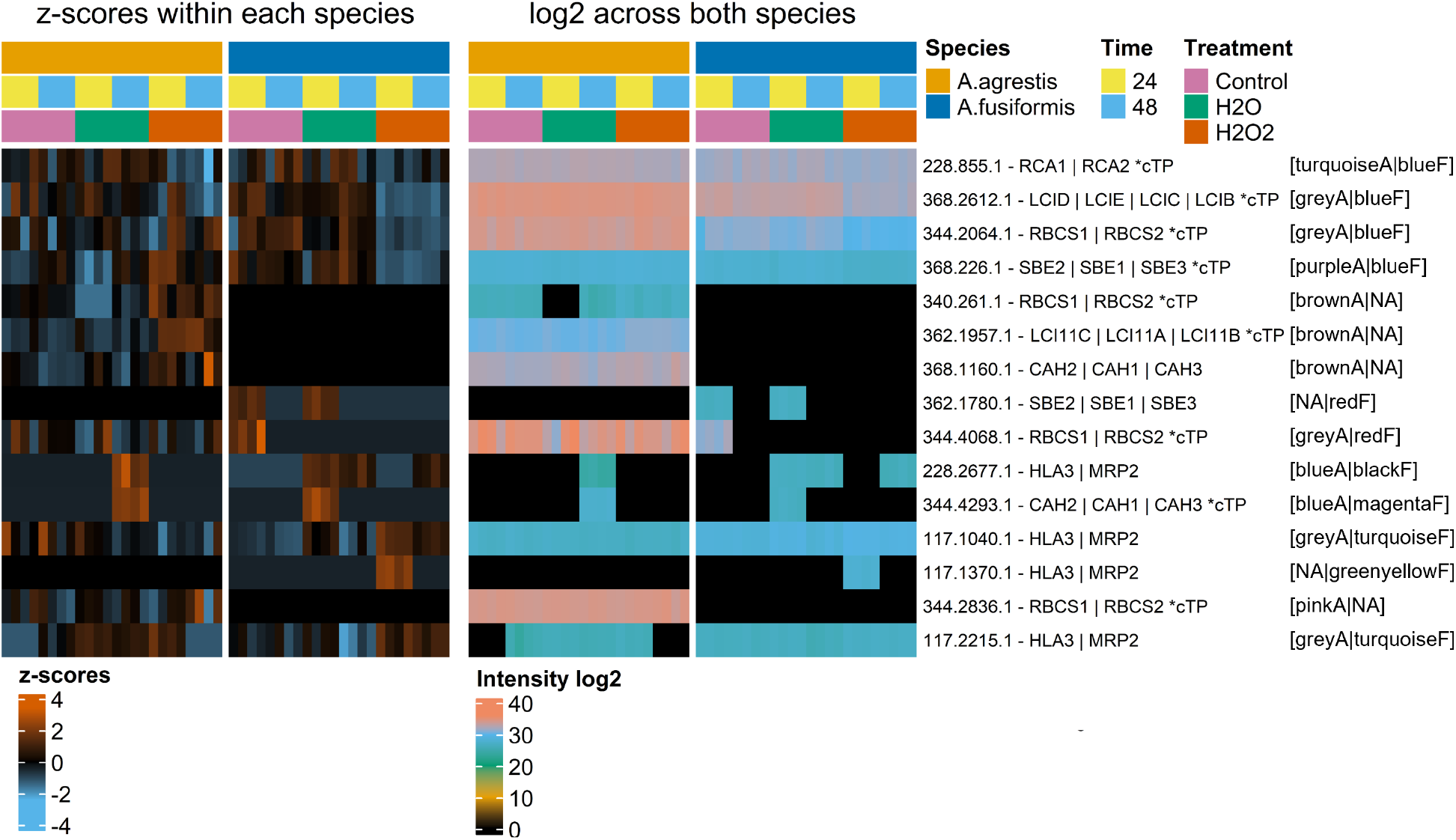
Heatmaps of core CCM-like proteins in *A. agrestis* and *A. fusiformis*: Intensities are shown in z-scores, calculated between all groups of each species (left), and shown in log_2_ across both species (right). The annotation above the heatmap indicates the time point, treatment, and species of each group. *A. agrestis* gene IDs are displayed with the corresponding homolog protein symbol of *C. reinhardtii*, and the presence of a predicted chloroplast transit peptide (cTP) is indicated by *cTP. The corresponding association of the detected core CCM-like proteins to the globally identified protein network cluster by WGCNA is listed after the protein description, starting with the module in *A. agrestis*, followed by the module in *A. fusiformis*. If the protein was not present in any module, the module color is replaced with “NA”.

### Homologs of CCM-involved proteins

Based on studies of *C. reinhardtii* proteins associated with CCM and pyrenoids, 912 proteins of interest were identified and categorized into Orthogroups (OGs) using computational methods (Supplemental Table S2). These proteins include those involved in photosynthesis, photorespiration, and other cellular processes connected to the CCM. Out of these 912 proteins, the location of 268 has been confirmed in *C. reinhardtii* (6–10). From the identified OGs, 768 possible homologs to 481 out of the initial 912 *C. reinhardtii* CCM-associated proteins were detected in *A. agrestis*, including a core set of essential CCM-associated proteins (see Supplemental Table S3) as well as 47 homologs to low CO_2_-inducible *C. reinhardtii* proteins. However, some CCM-associated proteins in *C. reinhardtii* do not have apparent homologs in *A. agrestis*, suggesting potential differences in the CCM process between the two organisms. Nonetheless, the presence of a core set of essential CCM-associated proteins in *A. agrestis* suggests that their functions may be conserved across species. Interestingly, seven of these CCM-associated homologs in *A. agrestis* (IDs: 344.3777, 368.1812, 368.2612, 228.5762, 340.1453, 228.5696, 228.5177) appear to be more similar to algal counterparts than to other land plants. Specifically, they cluster together with algal proteins rather than land plant proteins in the same OG. This group includes previously identified low CO_2_-inducible protein LCIB, a new candidate protein named LCI33 (Cre04.g217951), and several plastid-localized sulfate transporters such as SULP1 (Cre07.g348600) and SULP3 (Cre06.g257000). Additionally, there are three uncharacterized proteins in this category. LCI33 is a protein of unidentified function induced by low CO_2_ levels, whose expression does not change even if a pyrenoid is absent in *C. reinhardtii* (34). The close genetic relationship between LCI33 in *C. reinhardtii* and *A. agrestis* suggests that it might play a crucial role in the CCM function in hornworts, despite our failure to detect significant changes in its expression at the protein level.

## Discussion

Land plants have evolved various mechanisms to adapt to submersion. Here, we provide evidence that hornworts like *Anthoceros agrestis* can utilize a pyrenoid-based CCM to transition between submersive and atmospheric environments. In contrast, *A. fusiformis* responds to submersion with enhanced photorespiration and lipid droplet aggregation instead of carbon concentrating. This suggests that the hornwort CCM may aid in adaptation to flooding-desiccation cycles and maintaining optimal photosynthetic ability through a balanced O_2_/CO_2_ supply.

Submersion impacts land plants differently based on their habitat, lifestyle, and size. Algae in aquatic environments utilize active and passive HCO_3_^-^ import during CCM due to lower CO_2_ availability in water (27). Hornworts may adopt a similar mechanism when submerged, while terrestrial plants accessing atmospheric CO_2_ do not require such mechanisms. In 24-hour submersed *A. agrestis*, the total pyrenoid area increases due to a higher number of pyrenoids per plastid. This increase is accompanied by a greater number of stacked thylakoids associated with the pyrenoid region, potentially enhancing CO_2_ fixation efficiency. This induction of pyrenoids in *A. agrestis* during submersion indicates an adaptation to limited CO_2_ availability. This is supported by the upregulation of CCM-like proteins essential for importing and transforming inorganic carbon to CO_2_ for fixation, showing that hornworts possess effective CCM components. Consistent with this, several CCM-associated proteins are upregulated under submersion, including HLA3, an inorganic carbon transporter.

Both species showed similar Y(II) values under submersion, indicating comparable utilization of absorbed light for photochemistry. The slightly, though statistically non-significant, lower NPQ in *A. agrestis* underwater could be a result of efficient CO_2_ concentration near RuBisCO. The conversion of HCO_3_^−^ to CO_2_ + H_2_O in the lumen will take one H^+^ away from the proton-motive force and will make the lumen less acidic, which could explain the slightly lower NPQ in *A. agrestis*, as NPQ is pH-sensitive (35). In contrast, *A. fusiformis*’ higher NPQ might indicate a lower lumenal pH to dissipate excess energy more efficiently. This comparative analysis underscores the adaptation of *A. agrestis* to fluctuating aquatic-terrestrial environments through pyrenoid-based CCM.

The evolution of pyrenoids in hornworts was shown not to be correlated with atmospheric CO_2_ concentrations (13), which is currently debated (30). Submersion as an alternative explanation would partly match the respective species’ ecology. Both *A. agrestis* and *A. fusiformis* share a preference for moist, disturbed habitats, but they differ in their specific environmental tolerances and geographical distributions. *A. fusiformis* exhibits greater adaptability to a range of elevations and slightly drier conditions, whereas *A. agrestis* is more commonly associated with wetter and often flooded habitats in lowland environments (36). Therefore, sharing a pyrenoid-based CCM with algae may be advantageous for some hornworts, compared to others. Nonetheless, hornworts are land plants whose ancestors adapted to terrestrial environments, even though the latter may have been subject to periodic flooding.

Adjustments in pyrenoids occur only in the early stages of *A. agrestis*, within the first 24 hours of submersion in water, indicating a programmed response to submersion. By comparing molecular responses, plastid ultrastructural changes, and physiological performance during submersion and oxidative stress in *A. agrestis* and *A. fusiformis*, we identified specific regulation of CCM in both species. LC-MS/MS protein abundance analysis confirmed the presence of a core set of CCM-like proteins in both species under submersion and normal CO_2_ conditions. This suggests that pyrenoid-based CCM is constitutive in hornworts, especially *A. agrestis*, corroborated by findings that lowered atmospheric CO_2_ had little effect on pyrenoid dynamics and CCM-like gene expression (17).

Hornworts possess several algae-like homologs, including unique protein families that are exclusively shared with algae. Approximately half of the proteins involved in CCM of *C. reinhardtii* have homologs in *A. agrestis*, encompassing an essential core set of proteins, which suggests a similar functionality in CCM (6–10). These proteins in *A. agrestis* are differentially regulated, with some being upregulated under submersion or oxidative stress conditions. Submersion-induced core CCM-like proteins, such as cTP-containing CAH3 and LCI11, are homologous to *C. reinhardtii* counterparts. These types of proteins were reported to localize to plastids and, subcellularly, to pyrenoid-analogous locations in *A. agrestis* (17, 18). TEM revealed increased pyrenoid-like structures in *A. agrestis* within 24 hours of submersion (Figure 3), suggesting an algae-like, pyrenoid-based CCM underwater. Despite lacking pyrenoids, *A. fusiformis* upregulated core CCM-like proteins during submersion. Therefore, it remains unclear if *A. fusiformis* uses an alternative CCM, indicated by protein expression patterns under submersion and oxidative stress. In contrast, *A. fusiformi*s downregulates most core CCM-like proteins during H_2_O_2_ treatment, and Gene Ontology (GO) annotation analysis indicates higher photorespiration activity under these conditions, in line with our PAM measurements (Figure 5, Supplemental Table S8, and Supplemental Table S9).

Our findings suggest that oxidative stress induces a ROS-signaling cascade in *A. agrestis* for a subset of CCM-involved proteins (Supplemental Figure S9), excluding those related to pyrenoid formation. The focus here is on SF9 Cluster A2, which contains proteins upregulated under hyperoxia, which are not homologous to proteins directly involved in pyrenoid formation. This contrasts with the hyperoxia-induced pyrenoid formation observed in *C. reinhardtii* (29). The induction of CCM in hornworts appears to require different factors than in aquatic algae. Submersion may trigger a specialized signaling pathway in *A. agrestis* for pyrenoid reorganization and upregulation of CCM-involved proteins, partially overlapping with hyperoxia-inducible proteins. Conversely, the pyrenoid-absent *A. fusiformis* exhibits a divergent expression pattern for a small subset of CCM-related proteins between submerged and non-submerged conditions, indicating the absence of a submersion-based signaling pathway for CCM-related proteins in *A. fusiformis*.

Most *in silico*-identified CCM-like protein homologs in *A. agrestis* are also found in other land plant lineages. However, amino acid sequence similarities do not necessarily support the same functional roles. For instance, the luminal carbonic anhydrase CAH3 is unique to algae and typically absent in land plants, yet one of the CAH3 homologs in *A. agrestis* is predicted to have a chloroplast transit peptide, indicating a potentially similar function to that in algae (2). Co-expression of core CCM proteins, such as luminal CAH3 and thylakoid-channel-based LCI11, is crucial in *C. reinhardtii* for importing HCO_3_^-^ into the thylakoid lumen and converting it to CO_2_. The molecular basis for this process likely exists in *A. agrestis*, too, as evidenced by the presence of LCI11 homologs in its proteomic data, whereas LCI11 is absent in *A. fusiformis*. Notably, *A. agrestis* exhibited a significant increase in core CCM-like proteins in response to oxidative stress, a trend not observed in *A. fusiformis*. GO annotation analysis revealed an increased abundance of CO_2_-fixation and photosynthesis-related proteins in *A. agrestis* during hyperoxia. No pyrenoid-like structures were observed in *A. fusiformis* under any treatment by TEM analysis (Figure 3), suggesting that the expression of CCM-involved proteins like LCI11 might be linked to an existing pyrenoid-like structure.

The function and molecular basis of the complete CCM in hornworts remain unclear, and detailed similarities and differences between land-plant and algae-like pyrenoid-based CCMs still need to be elucidated. Exploration of functional domains in CCM-like protein sequences of *A. agrestis* supports the function of the corresponding *C. reinhardtii* proteins, although the involvement and process details in hornworts remain unknown. Identifying core CCM-like proteins in *A. agrestis* paves the way for uncovering the underlying mechanisms of hornworts’ CCM. Interactions among these proteins and additional necessary components, such as a linker protein for the pyrenoid matrix, remain to be found in hornworts (2). Considering that not all CCM-like proteins can be identified by homology, experimental testing for protein interactions could reveal new candidates for hornwort CCM proteins, as shown in *C. reinhardtii* (10). The presence of a functional pyrenoid in *A. agrestis*, but not in *A. fusiformis* underlines the need for further research to understand the genetic and molecular differences that govern CCM in hornworts.

The factors driving the sporadic evolutionary pattern of pyrenoid retention and loss in hornworts remain enigmatic. Adapting to life on land requires different strategies for water use, storage, and access compared to algae. Submersion poses a significant challenge for gametophytic plants with limited vertical growth, along with intermittent water availability. Our study shows that submersion triggers distinct abiotic responses in hornworts, varying between pyrenoid-present and pyrenoid-absent species. It is important to remember, however, that pyrenoids enhance photosynthetic efficiency by increasing carbon fixation and reducing photorespiration, both of which are critical for water-use efficiency (WUE).

Hornworts absorb water directly through their tissues and rely on diffusion for gas exchange (37). In habitats where (fresh) water availability is critical, enhanced photosynthetic efficiency aids in water conservation. By improving photosynthetic efficiency through the pyrenoid-based CCM, hornworts can grow and reproduce effectively with the water they absorb, achieving their growth and reproductive goals using less energy, leading to better survival and reproduction rates. Reducing the oxygenation reaction of RuBisCO through pyrenoids requires less energy and resources, sustaining growth and metabolic functions with less water. The oxygenation reaction produces 3-phosphoglycerate (3-PGA) and 2-phosphoglycolate (2-PG). Processing 2-PG involves multiple organelles (chloroplasts, peroxisomes, and mitochondria), releasing CO_2_, ammonia (NH_3_), and 3-PGA, while consuming additional ATP and NADPH. For each molecule of oxygen fixed by RuBisCO, an extra three ATP and two NADPH molecules are required beyond the standard Calvin cycle. To meet this demand, more light reactions are needed, where water is split by the oxygen-evolving complex of photosystem II to produce ATP and NADPH. Therefore, more efficient water use due to pyrenoids could contribute another explanation towards unraveling the ecological drivers of pyrenoid retention in some lineages.

Understanding the factors that induce pyrenoid formation and CCM in hornworts can provide insights into their eco-evolutionary significance and the water-to-land transition of plants. Efficient photosynthesis and suppression of RuBisCO’s oxygenation enable hornworts to thrive under both limited and excess water conditions. Submersion in *A. agrestis* induces a denser accumulation and increased number of pyrenoids, similar to changes observed under cold treatment (38).

Last but not least, our study revealed that *A. fusiformis* fills its open stroma spaces with lipid droplets, unlike the pyrenoid-forming *A. agrestis*. Under oxidative stress, notable shifts in CCM-like protein expression occur in both species without resulting in pyrenoid formation. Instead, hyperoxic conditions trigger plastid remodeling, resulting in various phenotypes such as small plastids, large starch-filled plastids, and plastids with irregular thylakoid arrangements (Figure 3). In hornworts, lipid droplets sometimes surround the pyrenoid matrix or are located in stroma spaces within the matrix (15, 16). Here in *A. fusiformis*, these plastid lipid droplet arrangements resemble the pyrenoid areas of *A. agrestis*, which have been described as “pyrenoid-like areas” ((15), p.123). Lipid droplets (LDs) serve various functions, including storage of lipids, enzymes, and substances, and accumulation of triacylglycerols (TAGs) for fats and oils. They are associated with the thylakoid membrane and can increase in size and number under stress (39). In *A. agrestis*, LDs are mostly present in chains or as single droplets across the plastid, while *A. fusiformis* shows significant LD accumulation even under unstressed conditions (Figure 3). In mutants of the model alga *Chlamydomonas reinhardtii* lacking starch under nitrogen starvation revealed an increase in TAG-producing enzymes, though lipid metabolism remained similar between mutants and the wildtype controls. Typically, *C. reinhardtii* stores carbon as starch first, followed by TAG accumulation; after starvation, starch is degraded before TAGs (40). Therefore, the higher abundance of LDs in *A. fusiformis* may be due to the absence of a pyrenoid-based CCM. Without effective carbon fixation like in *A. agrestis, A. fusiformis* may compensate by accumulating LDs as an alternative energy storage (41). This phenomenon was also observed in the charophyte alga *Zygogonium ericetorum*, where LDs fill open stroma spaces in the absence or dissolution of pyrenoids in desiccated algae (42). This indicates that LD presence and formation could be related to pyrenoid presence or absence, functioning as an additional energy storage form alongside starch. Puzzling in *A. fusiformis* are the expression patterns of proteins under submersion, which are similar to the control group, except for some core CCM-like proteins that are upregulated during the first 24 hours of submersion. This suggests that the co-expression of core CCM-like proteins may be controlled by the presence of a pyrenoid-like structure or might be genetically absent in *A. fusiformis*, warranting further investigation.

## Materials and Methods

### Plant material and experimental design

To study CCM in hornworts and identify CCM-relevant proteins, we compared the close relatives *Anthoceros agrestis*, which forms pyrenoids, and *A. fusiformis*, which is unable to form pyrenoids (15). Axenic cultures of both species were grown on KNOP medium (43) under ambient temperatures and light conditions. Two weeks before the experiments, axenic gametophytes were singled out into 6-well plates on KNOP medium, at a 14:10 day-night rhythm, with 23°C and 800 LUX at daytime and 21°C at night in the dark. Sample collection was done for the following two treatments: submersion with 3 ml autoclaved water, submersion with 3 ml autoclaved water and additional 100 µM hydrogen peroxide, to imitate a hyperoxic environment (29); as controls, we used plants in 6-well plates in atmospheric, non-submersion conditions. Sampling time points were 24h and 48h with four replicates each and per treatment.

### Mass spectrometry

For the sample preparation, the whole gametophyte was harvested, frozen in liquid nitrogen, and pulverized. Extraction was done in lysis buffer and after FASP method (44). Protein concentration was measured with the Pierce TM BCA Protein Assay kit (Thermo Scientific, Waltham, MA, USA). 100 µg of proteins were digested by Trypsin enzyme (PROMEGA, Madison, WI, USA). For desalting of peptide aliquots, Stage Tips C18 were used (45) and afterward dried by vacuum centrifugation (Concentrator Plus, Eppendorf) and frozen for storage. The peptide pellet was resuspended in LC running buffer (2% (v/v) acetonitrile/0.05%(v/v) trifluoroacetic acid in Millipore water) at a theoretical concentration of 1 mg/mL peptide mixture and separated with an Ultimate 3000 RSLCnano System (Thermo Scientific) on a separation column (Acclaim PepMap100 C18, 75 mm i.D., 2 mm particle size, 100 Å pore size; Thermo Scientific) with a length of 50 cm. The LC system was coupled with a nanospray source to a Q Exactive Plus mass spectrometer (Thermo Scientific) while the process was monitored over Xcalibur (Thermo Scientific).

### Label-free proteomic data analysis

Raw data analysis was performed with MSFragger using Fragpipe (46–49), and for both the false discovery rate (FDR) was set to 1%. A protein database for *A. agrestis* (https://www.hornworts.uzh.ch/en/download.html, UZH Switzerland) was used for protein identification in both species since no *A. fusiformis* database was available at that time point. Translated proteome sequences based on the newly released genome of *A. fusiformis* (available at hornwortbase.org since January 2025, i.e., after our analyses were completed) were subsequently queried using BLASTP for the core CCM proteins identified in *A. agrestis*. Homologs for all *A. agrestis* core CCM proteins were detected in *A. fusiformis*. Notably, the protein domains were conserved between the two species, although the extent of sequence similarity varied. These findings suggest that reanalyzing our data with the updated genome would likely not result in substantially improved protein identification. The database was expanded in protein descriptions by using *InterProScan* (50) and *EggNOG* v5.0 (51). The *MaxLFQ Intensity* results of the *combined_peptide*.*tsv* output of Fragpipe were used for further data analysis. The data was cleaned up by removing samples with no or very low recorded intensities. The cleaned-up data was further analyzed in R (https://www.R-project.org/): The principal component analysis (PCA) was performed with the *factoextra* package (https://CRAN.R-project.org/package=factoextra). Pair-wise differences in protein abundance between treatments were calculated by *DEqMS* (52) and visualized with *EnhancedVolcano* (https://github.com/kevinblighe/EnhancedVolcano). In addition, a subset, which included only proteins that were not present in the control treatments, was used for visualization of intersections between the different submersion treatments with *UpSet* (53), followed by a GO-Term overrepresentation of the intersection with the *enricher* function of *clusterProfiler* (54).

### Identification of protein clusters

To identify groups of proteins with the same expression pattern between treatments, weighted correlation network analysis was performed with *WGCNA* (55), for each species separately, using the *blockwiseModules* function with *networkType* set to “signed” and *corType* set to “bicor”. In addition, the kinetic pattern of significantly differentially expressed proteins was calculated by *MaSigPro* (56), and cluster number was chosen by K-means cluster analysis of the significantly expressed protein subset.

### *In silico* analysis of gene families

We compiled a dataset of 30 readily available plant proteomes (Supplemental Table S4), ranging from Chlorophytes and Charophytes, over non-vascular land plants to different genera of vascular land plants. We subjected these data to an *OrthoFinder2* analysis (57), to identify Orthogroups (OGs) of ortholog proteins between species. The resulting data were post-analyzed with TargetP 2.0 (58) to identify proteins with transit peptides, directing to the chloroplast (cTP) or lumen (luTPs), herein combined and referred to as cTPs. OGs containing CCM-involved proteins from *C. reinhardtii* (6–10) were selected and subjected to a second filtering, where we searched for the presence of *A. agrestis* protein homologs in OGs with CCM proteins of *C. reinhardtii*. Those candidate CCM-like proteins of *A. agrestis* were double-checked with *InterProScan* to infer protein functions in the form of functional identified protein domains and compared to the leading counterpart CCM-protein of *C. reinhardtii*.

### Transmission Electron Microscopy (TEM)

Hornwort material was prefixed in 4% glutaraldehyde adjusted with a 0.1 M sodium cacodylate buffer at a pH of 7.2 and subsequently washed in the same buffer. Post-fixation was done in 1% osmium tetroxide dissolved in the same buffer as used for prefixing. Afterward, the specimens were washed in ddH_2_O and dehydrated in a graded series of ethanol, followed by propylene oxide. Embedding was done with Spurr low-viscosity embedding medium (Electron Microscopy Sciences, Hatfield, PA, USA) following the manufacturer’s instructions. 60nm sections were produced with a Reichert Ultracut S ultramicrotome (Leica, Vienna, Austria) using an ultra 45° diamond knife (DiATOM, Nidau, Switzerland). Slices were placed either on uncoated hexagonal mesh or on Formvar-filmed slot TEM copper grids (Plano, Wetzlar, Germany) and, thereafter, stained with uranyl acetate and Reynolds lead citrate using a Phoenix contrasting machine (Staining Technologies, Berlin, Germany). Imaging was done on an EM 900 TEM (Zeiss, Oberkochen, Germany) equipped with a 2K wide-angle slow-scan CCD camera (Tröndle Restlichtverstärkersysteme, Moorenweis, Germany). Measurements were made with Fiji (https://imagej.net/software/fiji/) for multiple samples per treatment and tested with data suiting statistical analysis for significance in time- and treatment-related differences between groups on multiple variables.

### Pulse Amplitude Modulation (PAM) measurements

The experiment was conducted with three biological replicates for the submersion in H_2_O and H_2_O_2_, and the samples were randomly arranged in 6-well plates. The measuring points were taken at 0, 24, and 48 hours of submersion as well as during recovery of 24 hour after submersion. The chlorophyll fluorescence was measured using a Maxi-Imaging Pulse-Amplitude-Modulation (PAM) chlorophyll fluorometer (Walz, Germany). After 30 minutes of dark incubation, the plants were exposed to a photosynthetically active radiation intensity of 285 PPFD for 10 minutes, which we determined through previous light curve analysis as the optimal level for both *A. agrestis* and *A. fusiformis* (Supplemental Figure S10). This was followed by a 10-minute measurement in the dark, monitored by the ImagingWin software (Walz). Before the measurements, the lid was removed. The PSII operating efficiency was calculated as Y(II) = (Fm’-F)/Fm’, and the non-photochemical quenching of PSII fluorescence was calculated as NPQ = (Fm-Fm’)/Fm’.

## Acknowledgements

We thank Peer Martin (HU Berlin) and Dr. Martin Scholz (University of Münster) for excellent technical assistance during TEM and MS sample preparation, respectively. This project received support from the Deutsche Forschungsgemeinschaft (joint DFG grant: HI739/20-1/SZ368/2-1/WI4507/9-1 to M.H., P.S., and S.W.) as part of the DFG-funded priority program *MAdLand* SPP2237 (https://madland.science).

## Notes

**Competing Interest Statement:** The authors declare no conflict of interest.

### Competing Interest Statement

The authors have declared no competing interest.

