## Supplemental for "Submersion and oxidative stress triggers pyrenoid formation, carbon-concentration-related protein remodeling and sub-plastidial rearrangements in hornworts": Supplemental_Figures.Noetzold_etal.pdf

[1] Institute of Plant Biology and Biotechnology, University of Münster, Münster 48149, Germany; [2] Department of Plant Evolution and Biodiversity, Institute for Biology, Humboldt-University Berlin, Berlin 10115, Germany; [3] Department of Molecular Parasitology, Institute for Biology, Humboldt-University Berlin, Berlin 10115, Germany; [4] Department of Systematic and Evolutionary Botany, University of Zurich, Zurich CH-8006, Switzerland and Zurich-Basel Plant Science Center, ETH Zurich, Zurich CH-8092, Switzerland; [5] Institute of Plant Science and Resources, Okayama University, Kurashiki 710-0046, Japan; [6] Institute for Evolution and Biodiversity, University of Muenster, Muenster 48149, Germany

* Correspondence:
Susann Wicke and Michael Hippler

**This PDF includes Figures S1 to S10**

**Supplemental Tables S1 to S10 available for download in separate files.**

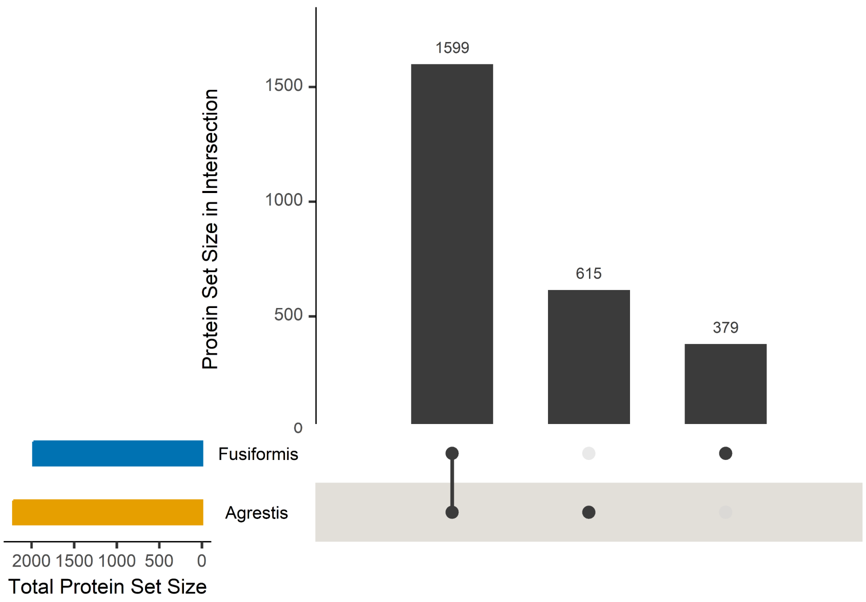


**Supplementary Figure 1: UpSet plots for the identified proteins in *A. agrestis* and *A. fusiformis* tissue.** Protein set intersections are in descending order with the total size given by the number above each bar. Filled circles show intersections and connection trough lines indicate intersection groups. The intersections show the number of proteins found present in both species (n=1599), or separately in *A. agrestis* (n=615) or *A. fusiformis* (n=379).

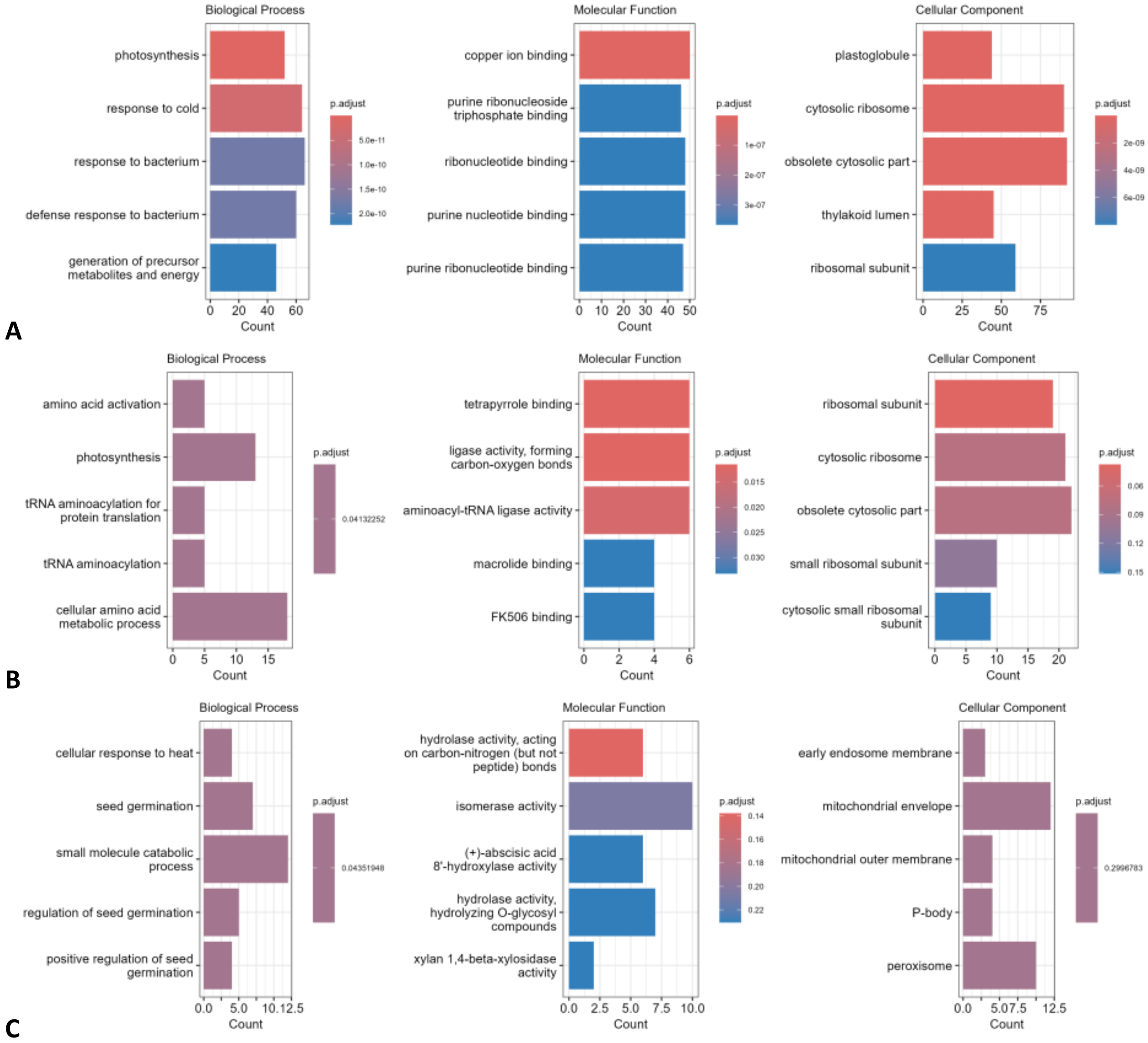


**Supplementary Figure 2: GO-term overrepresentations of protein intersection between *A. agrestis* and *A. fusiformis*, based on the UpSet plot for all identified proteins within both species.** GO-ORA was separated into the three main GO-term groups: Biological process, molecular function, and cellular component, showing the Top 5 GO-terms. (A) GO-ORA for 1599 proteins expressed in both hornwort species. (B) GO-ORA for 615 proteins only expressed in *A. agrestis*. (C) GO-ORA for 379 proteins only expressed in *A. fusiformis*. For further information see Supplemental Figure 1.

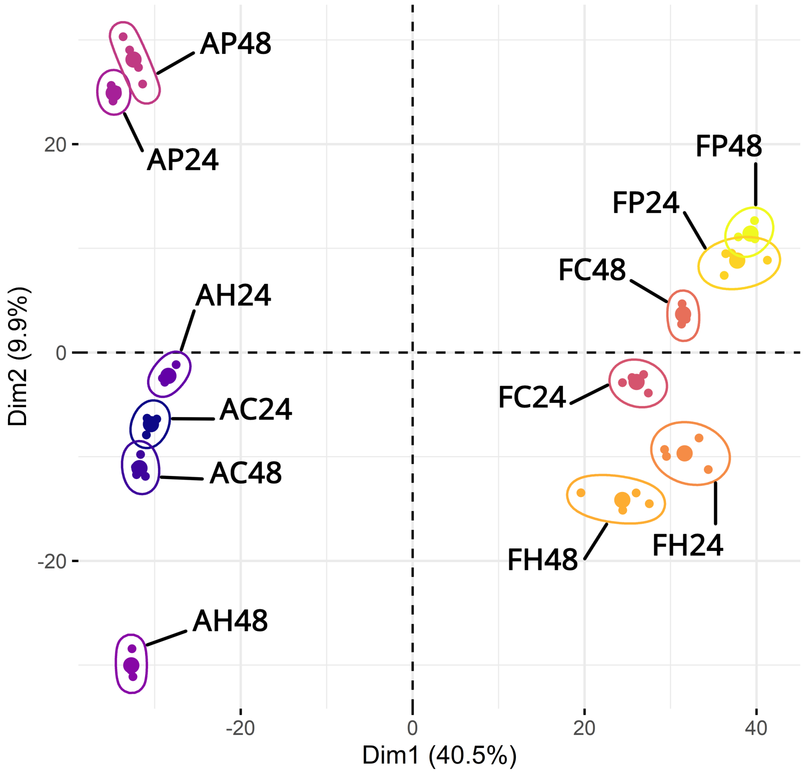


**Supplementary Figure 3: Principal component analysis of LC-MS/MS samples of *A. agrestis* and *A. fusiformis*.** 48 LC-MS/MS samples are grouped by treatment and time point: Non-submersed 24h (C24), non-submersed 48h (C48), $H_2O$ submersion 24h (H24), $H_2O$ submersion 48h (H48), $H_2O_2$ submersion 24h (P24), and $H_2O_2$ submersion 48h (P48). Species are indicated by A (*A. agrestis*) and F (*A. fusiformis*). Dim1 shows the PCA component 1 (40.5%), while Dim2 shows the PCA component 2 (9.9%), combined explaining 50.4 % of the variation within the measured intensities for all samples.

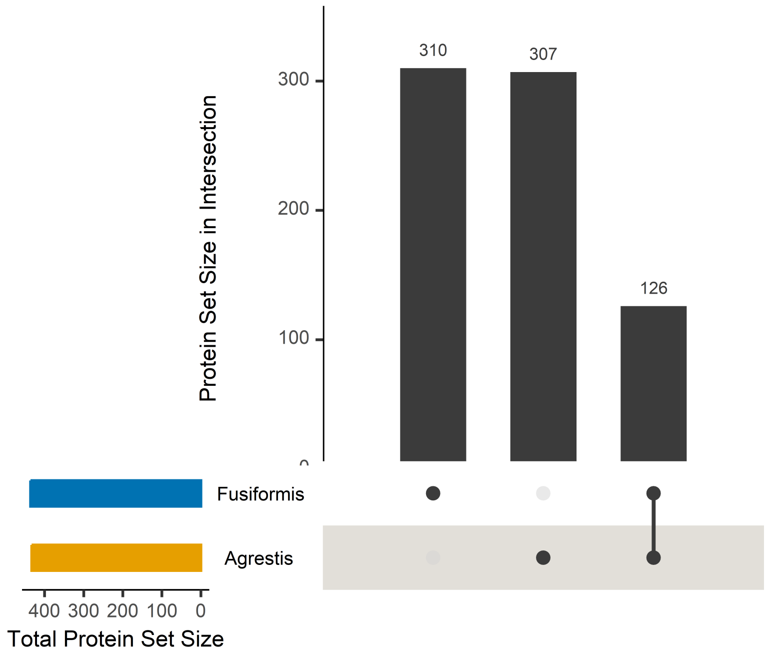


**Supplementary Figure 4: UpSet plots for the identified and significantly expressed proteins in *A. agrestis* and *A. fusiformis* tissue.** Protein set intersections are in descending order with the total size given by the number above each bar. Filled circles show intersections and connection trough lines indicate intersection groups. A total of 126 proteins were shared between both species, expressed in all treatments, but significantly differently regulated, while *A. fusiformis* showed 310 individuals significantly expressed proteins and *A. agrestis* a total of 307 proteins.

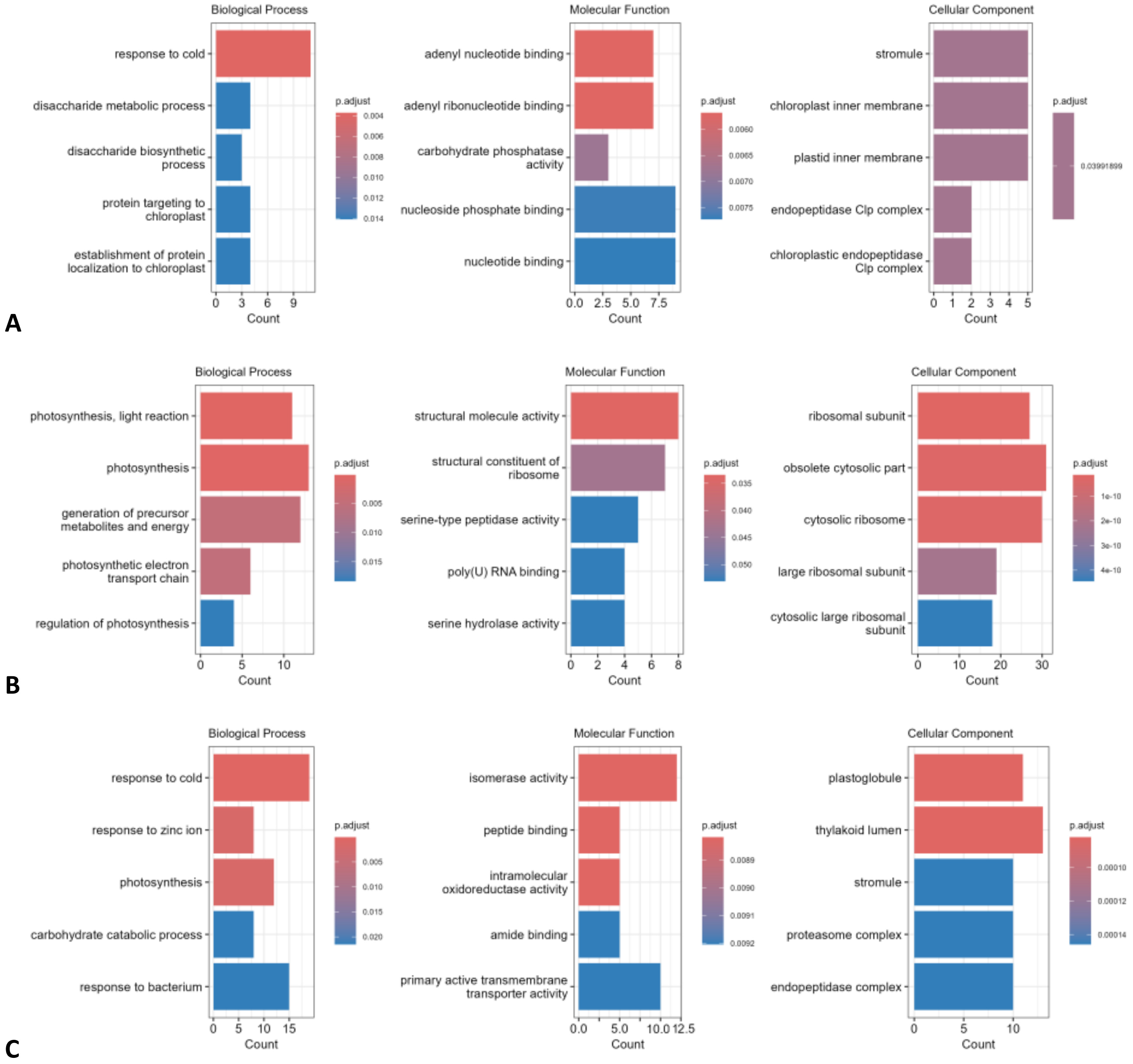


**Supplementary Figure 5: GO-term overrepresentations of protein intersection between significantly expressed proteins in *A. agrestis* and *A. fusiformis*, based on the UpSet plot for the significantly expressed identified proteins within both species.** GO-ORA was separated into the three main GO-term groups: Biological process, molecular function, and cellular component, showing the Top 5 GO-terms. (A) GO-ORA for 126 proteins expressed in both hornwort species. (B) GO-ORA for 307 proteins only expressed in *A. agrestis*. (C) GO-ORA for 310 proteins only expressed in *A. fusiformis*. For further information see Supplementary Figure 4.

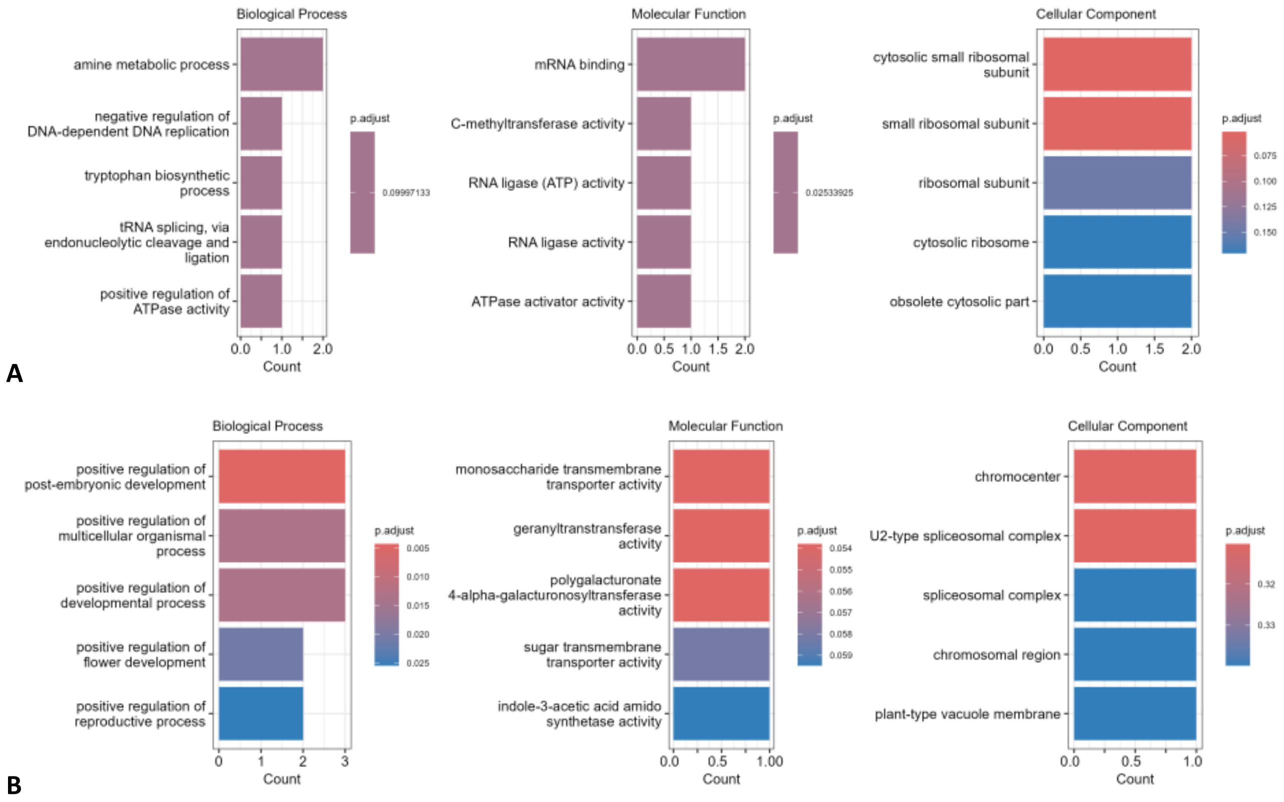


**Supplementary Figure 6: GO-term overrepresentations of the Top 20 significantly expressed proteins in *A. agrestis* and *A. fusiformis*.** GO-ORA was separated into the three main GO-term groups: Biological process, molecular function, and cellular component. (A) Top 20 significant proteins in exclusively *A. agrestis* identified proteins. (B) Top 20 significant proteins in exclusively *A. fusiformis* identified proteins.

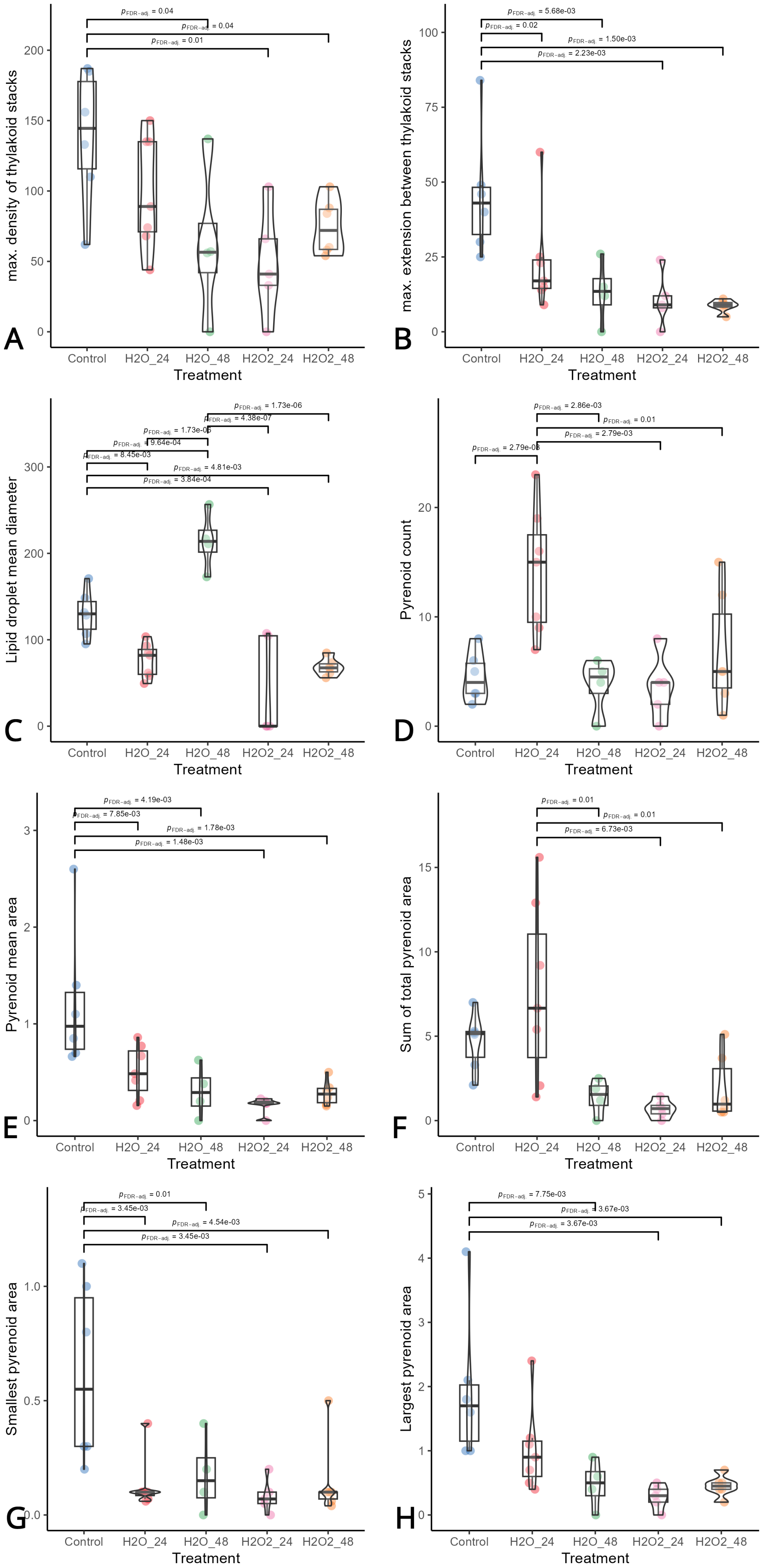

A
max. density of thylakoid stacks
pFDR-adj. = 0.04
pFDR-adj. = 0.04
pFDR-adj. = 0.01
B
max. extension between thylakoid stacks
pFDR-adj. = 5.68e-03
pFDR-adj. = 0.02
pFDR-adj. = 1.50e-03
pFDR-adj. = 2.23e-03
C
Lipid droplet mean diameter
pFDR-adj. = 1.73e-06
pFDR-adj. = 4.38e-07
pFDR-adj. = 1.73e-05
pFDR-adj. = 9.64e-04
pFDR-adj. = 8.45e-03
pFDR-adj. = 4.81e-03
pFDR-adj. = 3.84e-04
D
Pyrenoid count
pFDR-adj. = 2.86e-03
pFDR-adj. = 0.01
pFDR-adj. = 2.79e-03
pFDR-adj. = 2.79e-03
E
Pyrenoid mean area
pFDR-adj. = 4.19e-03
pFDR-adj. = 7.85e-03
pFDR-adj. = 1.78e-03
pFDR-adj. = 1.48e-03
F
Sum of total pyrenoid area
pFDR-adj. = 0.01
pFDR-adj. = 0.01
pFDR-adj. = 6.73e-03
G
Smallest pyrenoid area
pFDR-adj. = 0.01
pFDR-adj. = 3.45e-03
pFDR-adj. = 4.54e-03
pFDR-adj. = 3.45e-03
H
Largest pyrenoid area
pFDR-adj. = 7.75e-03
pFDR-adj. = 3.67e-03
pFDR-adj. = 3.67e-03
Control
H2O_24
H2O_48
H2O2_24
H2O2_48
Treatment

**Supplementary Figure 7: Boxplots with statistical information on the comparison of the transmission electron microscopy measurements between the different submersion treatments in *A. agrestis*.** Figures were created with the R package ggstatsplot (Patil, 2021) and tested for statistical significance with one-way ANOVA. Significant differences between treatments are indicated above the boxplots. Violin plots indicate data distribution, while colored points display the individual data points. Treatment included the non-submersed control, $H_2O$ submersion after 24h and 48h, and $H_2O_2$ submersion after 24h and 48h. **(A)** Boxplot of the maximal density of stacked thylakoid between pyrenoid, within the overall pyrenoid area within the plastid. **(B)** Boxplot of the maximal observed extension between thylakoid stacks between pyrenoids. **(C)** Boxplot of the lipid droplet mean diameter per plastid. **(D)** Boxplot of the total pyrenoid count per plastid. **(E)** Boxplot of the mean pyrenoid area per plastid, calculated as follows: sum of all pyrenoid areas per plastid / total count of pyrenoid per plastid **(F)** Boxplot of the sum of the total pyrenoid area per plastid. **(G)** Boxplot of the area of the smallest pyrenoids per plastid. **(H)** Boxplot of the area of the largest pyrenoids per plastid.

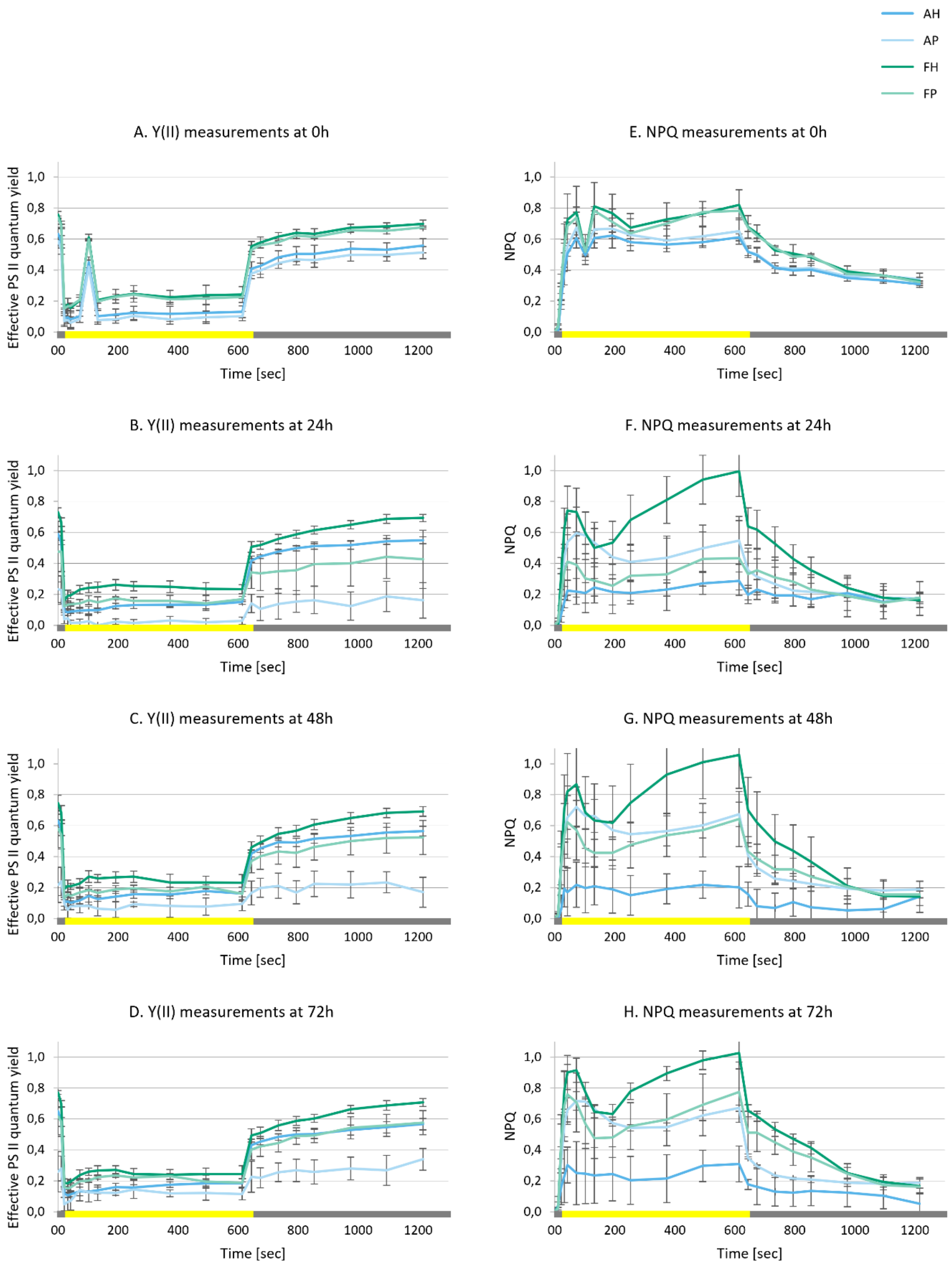


**Supplementary Figure 8. Effective PS II quantum yield (Y(II)) and non-photochemical quenching (NPQ) for submersed hornworts.** Figures A to D show the Y(II) for $H_2O$ (H) and $H_2O_2$ (P) submersed *A. agrestis* (A) and *A. fusiformis* (F). Figures E to H show the NPQ measurements. The x-axis gives the time frame of experimental measurements and the values for light (yellow bar) and dark (grey bar) treatment over the time frame of 20 minutes. The standard deviation of the averaged 3-replicate values is given in the error bars.

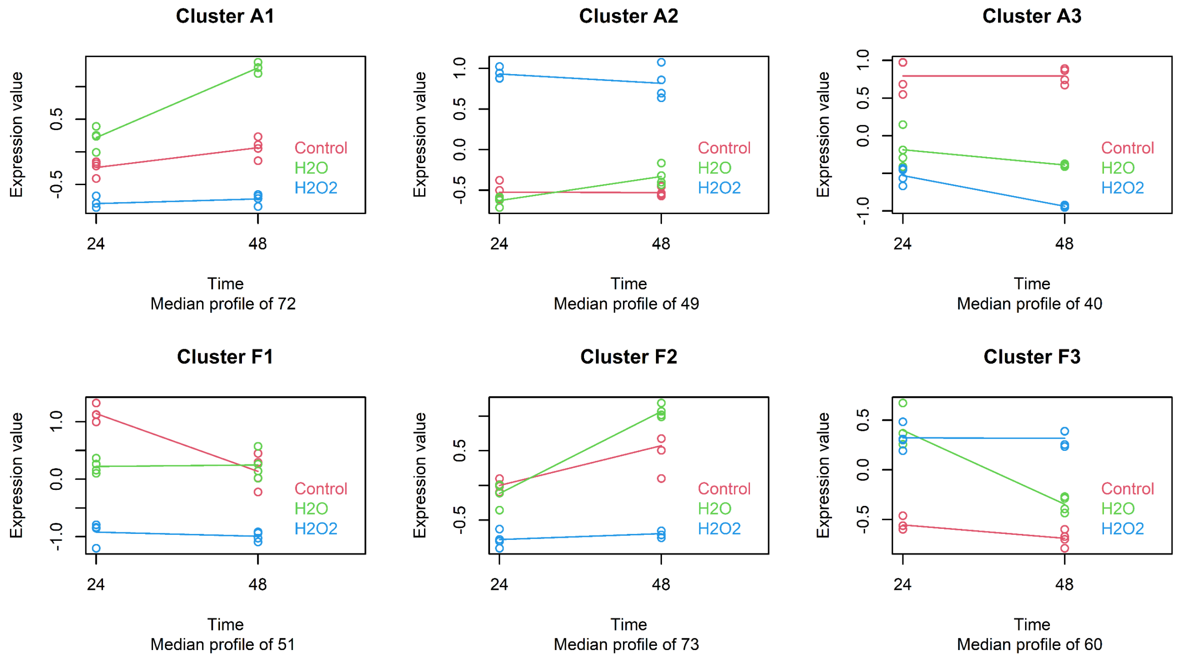


**Supplementary Figure 9: Kinetic clusters of significantly differently expressed CCM-related proteins in *A. agrestis* and *A. fusiformis*.** Differential expression analysis was performed with MaSigPro and k-clustering method. Top row: Three clusters were identified for a total of 161 proteins in *A. agrestis* (A1, A2, and A3). Bottom row: Three clusters were identified for 184 proteins in *A. fusiformis* (F1, F2, and F3). The three treatment groups control (red), $H_2O$ (green), and $H_2O_2$ (blue) are displayed over the time frame of 24 to 48 hours.

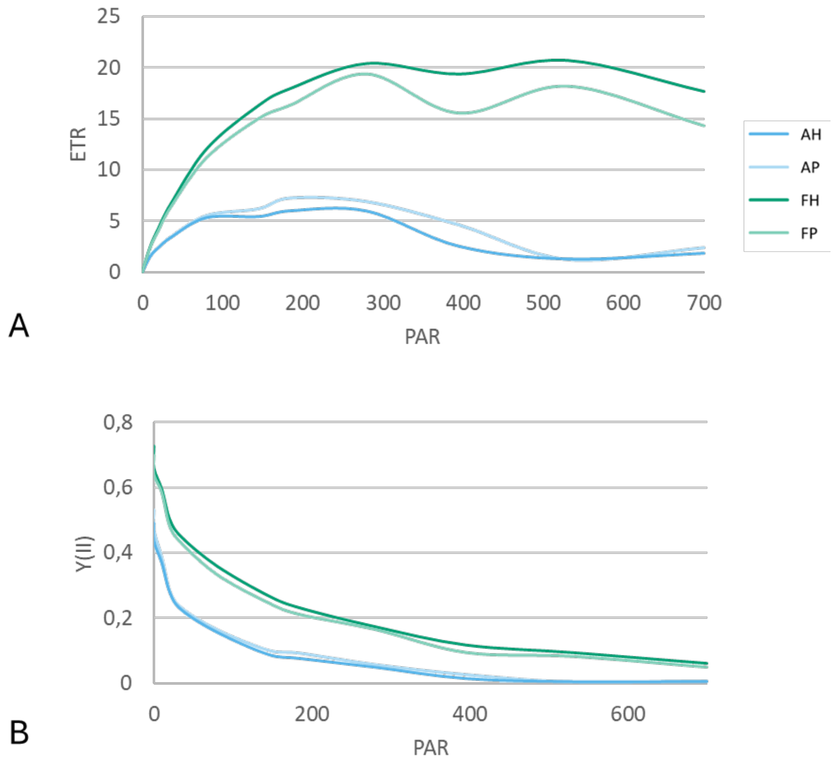


**Supplementary Figure 10: Light curve measurements of *A. agrestis* and *A. fusiformis*.** The duration of light intervals was set to 30s with previous acclimation in the dark for 30 min for *A. agrestis* and *A. fusiformis*. (A) Light electron transport rate (ETR) curve, with ETR calculated using the photosystem II yield (YII) data. (B) Light Y(II) curve ranges from 1 (100%) to 0 (0%) of absorbed light energy directed to PSII.
